# Mechanism of ZNF143 in chromatin looping and 3D genome organization

**DOI:** 10.1101/2023.09.27.559875

**Authors:** Mo Zhang, Haiyan Huang, Jingwei Li, Qiang Wu

## Abstract

The transcription factor ZNF143 contains a central domain of seven zinc fingers in a tandem array and is involved in 3D genome construction; however, the mechanism by which ZNF143 functions in chromatin looping remains unclear. Here, we show that ZNF143 directionally recognizes a diverse range of genomic sites within enhancers and promoters and is required for chromatin looping between these sites. In addition, ZNF143 is located between CTCF and cohesin at numerous CTCF sites, and ZNF143 removal narrows the space between CTCF and cohesin. Moreover, genetic deletion of ZNF143, in conjunction with acute CTCF depletion, reveals that ZNF143 and CTCF collaborate to regulate higher-order chromatin organization. Thus, ZNF143 is recruited by CTCF to the CTCF sites to regulate TAD formation, whereas directional recognition of genomic DNA motifs directly by ZNF143 itself regulates promoter activity via chromatin looping.

## Introduction

CTCF is a key mammalian architectural protein for interphase 3D genome folding^1,2^. Specifically, CTCF dynamically recognizes a wide range of genomic sites within the linear 1D sequences known as CBS (CTCF binding site) elements. Their genomic distributions and relative orientations determine the looping specificity of long-distance chromatin interactions^1,3–6^. In particular, there is a strong tendency for close spatial contacts between forward-reverse convergent CBS elements^7–10^. Mechanistically, asymmetrically stalling of loop-extruding cohesin complexes at convergent CTCF sites results in their close spatial contacts^11,12^ since orientated CTCF interacts with the cohesin complex via its N-terminus but not C-terminus^13–15^. These convergent tandem CBS elements were recently shown to function as insulators to balance topological promoter-enhancer selection and to block improper activation of non-cognate promoters by remote enhancers^16–19^.

As a paradigm to investigate mechanisms of genome folding, the human clustered protocadherin (*cPCDH*) genes are organized into three sequentially- linked clusters of *PCDH α*, *β*, and *γ*, spanning a region of ∼1 M bp (Figure 1A)^20–22^. This complex locus forms a superTAD comprising *PCDH α* and *βγ* subTADs, with 15 and 38 (*16 β* and *22 γ*) variable exons, respectively, followed by single sets of three downstream constant exons within each subTAD (Figure 1A). Specifically, the *PCDHα* subTAD contains a repertoire of 13 alternate (*α1*-*α13*) and 2 c-type (*αc1*- *αc2*) variable exons each preceded by a separate promoter. Each of the 13 alternate variable promoters is flanked by two forward-oriented CBS elements (CSE for conserved sequence element and eCBS for exonic CBS) and the *αc1*, but not *αc2*, promoter is associated with a single CBS element, resulting in a tandem array of 27 forward-oriented CBS elements. By contrast, the *HS5-1* enhancer element, which is located at the boundary between the *PCDH α* and *βγ* subTADs, is flanked by two reverse-oriented CBS elements (*HS5-1a* and *HS5-1b*)^7,16,23^. Continuous active “loop extrusion” by cohesin complexes anchored by these convergent forward-reverse CBS elements results in long-distance chromatin interactions between the *HS5-1* enhancer and target variable promoters. The activation of variable promoters by the *HS5-1* enhancer determines the stochastic and allelic *cPCDH* gene choice in single cells in the brain^10,16,24–26^.

**Figure 1.**
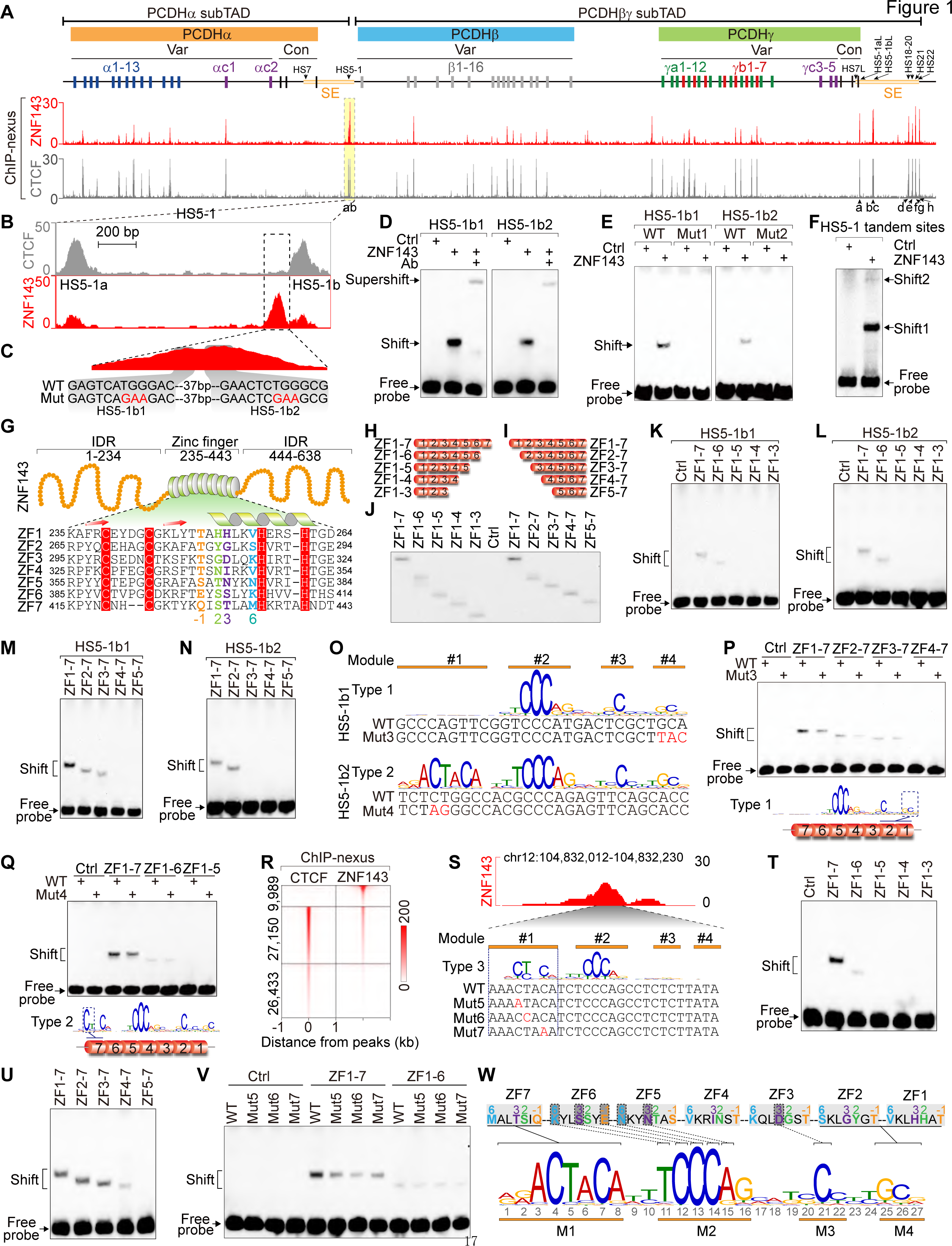
Directional ZNF143 recognition of tandem SBS elements within the *PCDH HS5-1* enhancer. (A) ZNF143 and CTCF ChIP-nexus profiles at the human clustered *PCDH* locus, showing co-localization of CTCF and ZNF143 at most variable promoters and super-enhancers. This locus comprises three tandem-arrayed *PCDH α*, *β*, and *γ* clusters. Both *PCDH α* and *γ* clusters contain multiple variable (Var) first exons, each of which is alternatively *cis*-spliced to a single set of cluster-specific constant (Con) exons. The *PCDHβ* cluster has only 16 variable exons, but with no constant exon. These three clusters form a superTAD with two (*α* and *βγ*) subTADs, each of which has its own super-enhancer (SE) at its downstream boundary. (B) Close-up of the *HS5-1* enhancer region showing the colocalization of ZNF143 and CTCF at the two CBS elements (*HS5-1a* and *HS5-1b*), as well as the unique binding of ZNF143 at a region immediately upstream of *HS5-1b*. (C) Enlargement of the unique ZNF143 peak reveals two subpeaks of 37 bp- spaced SBS elements of *HS5-1b1* and *HS5-1b2* in tandem. (D) Specific binding of the recombinant ZNF143 to the DNA probes of *HS5-b1* or *HS5-1b2* by the electrophoretic mobility shift assay (EMSA). (E) Mutation (Mut1 or Mut2) of *HS5-b1* or *HS5-1b2* abolishes the ZNF143 binding. (F) EMSA of ZNF143 binding to a DNA probe containing the two tandem SBS elements of *HS5-1b1* and *HS5-1b2,* generating two shifted bands. (G) Schematic of the ZNF143 central zinc finger (ZF) domain with the flanking N- and C-terminal intrinsically disordered regions (IDR). The amino acid sequences of the seven tandem ZFs are shown in the bottom. The residues at the positions -1, 2, 3, and 6 of ZF α-helix are highlighted in yellow, green, purple, and blue, respectively. The cysteine (C) and histidine (H) residues that contact with the zinc ion are highlighted in red background. (**H** and **I**) Schematics of the truncated ZNF143 proteins, with sequential deletions of ZFs from either C- (**H**) or N-terminus (**I**). (**J**) Western blot of recombinant ZNF143 proteins with sequential deletions of ZFs. (**K-N**) EMSA of truncated ZNF143 proteins with the *HS5-1b1* (**K** and **M**) or *HS5- 1b2* (**L** and **N**) SBS probe. (O) Sequences of the mutant probes (Mut3 and Mut4) of *HS5-1b1* and *HS5- 1b2*. (P) Effect of Mut3 on the binding of a series of N-terminal truncated ZNF143. (Q) Effect of Mut4 on the binding of a series of C-terminal truncated ZNF143. (R) CTCF and ZNF143 ChIP-nexus heatmaps of ZNF143 unique peaks, co- localized peaks, and CTCF unique peaks. (S) Sequences of three Module1 mutant probes (Mut5-7). The corresponding ZNF143 ChIP-nexus peak is shown above. (**T** and **U**) EMSA of ZNF143 with sequential deletions of ZFs from either C- (**T**) or N- (**U**) terminus. (U) Effect of Module1 mutation on EMSA of ZNF143. (V) Schematic of ZNF143 binding to its cognate SBS element.

The transcription factor zinc finger protein 143 (ZNF143) attracts increasing attention as a 3D genome modulator due to its ubiquitous expression and genome-wide co-localization with CTCF and cohesin^27–30^. ZNF143 is known for its ability to activate transcription of protein-coding and snRNA genes^31–34^. Specifically, ZNF143 is known as STAF for selenocysteine tRNA gene transcription-activating factor, and hence its genomic binding site is known as SBS (STAF binding site)^33,35^. Despite intensive research on ZNF143 in 3D genome organization^29,36,37^, its DNA recognition mechanism and role in higher-order chromatin organization remain unclear. Here we report that direct directional recognition of genomic sites by ZNF143 is required for long-distance chromatin contacts of promoters and that ZNF143 is crucial for genome compartmentalization.

## Results

### Directional ZNF143 recognition of SBS elements within PCDH HS5-1 enhancer

We performed ChIP-nexus (chromatin immunoprecipitation experiments with nucleotide resolution through exonuclease, unique barcode, and single ligation) experiments by using a specific antibody against ZNF143 and found that, similar to CTCF, ZNF143 is enriched at most *PCDH* variable promoters as well as super-enhancers (Figure 1A). Specifically, ZNF143 and CTCF are mostly colocalized at the *CSE* and *eCBS* elements in the variable promoter region of the *PCDHα* cluster (Figures 1A and S1A-S1T). However, in the *HS5-1* enhancer region, we found that, in addition to the colocalization of ZNF143 with CTCF at the two CBS elements (*HS5-1a* and *HS5-1b*), a single ZNF143 peak is localized immediately upstream of *HS5-1b* (Figure 1B). Enlargement of this single ZNF143 ChIP-nexus peak revealed two subpeaks, which contain two tandem SBS elements, termed *HS5-1b1* and *HS5-1b2*, in the reverse orientation separated by only 37 bp (Figure 1C). This suggests that ZNF143 separately recognizes each of these two tandem SBS elements.

We performed EMSA experiments using the *HS5-1b1* or *HS5-1b2* probe and confirmed the direct binding of ZNF143 to both SBS elements (Figure 1D). In addition, we generated two mutations, Mut1 and Mut2, and found that they abolish the ZNF143 binding (Figures 1C, 1E, and S1U). Finally, we observed two shifted bands using one EMSA probe containing both *HS5-1b1 and HS5- 1b2*, suggesting that each is recognized by a single ZNF143 protein (Figure 1F).

We conclude that ZNF143 binds directly to the *PCDHα HS5-1* enhancer which contains two tandem SBS elements each recruiting a single ZNF143 protein.

ZNF143 contains a central DNA-binding domain (DBD) with 7 tandem-arrayed C_2_H_2_-type zinc fingers (ZF1-7), which are flanked by N- and C-terminal intrinsically disordered regions (IDRs) (Figures 1G and S1V). To investigate the mechanism of ZNF143 binding to its cognate sites, we generated a series of truncated ZNF143 proteins through sequential deletions of ZFs from either C- or N- terminus (Figures 1H-1J). We performed comprehensive EMSA experiments with both *HS5-1b1* and *HS5-1b2* SBS probes (Figures 1K-1N). We found that C-terminal deletions up to ZF6 (ZF1-5) abolish the binding of ZNF143 to both SBS probes (Figures 1K and 1L). In addition, N-terminal deletions up to ZF3 (ZF4-7) or ZF2 (ZF3-7) abolish the binding to *HS5-1b1* or *HS5-1b2*, respectively (Figures 1M and 1N). This suggests that ZF3-6 or ZF2-6 is essential for ZNF143 to bind to *HS5-1b1* or *HS5-1b2*, respectively.

Interestingly, both SBS elements within the *HS5-1b* enhancer region are in the reverse orientation (We define that the direction of Module1-4 is the forward orientation) (Figure 1O). To investigate the directionality of ZNF143 binding, we generated two additional mutant probes, Mut3 and Mut4, within the Module4 of *HS5-1b1* and Module1 of *HS5-1b2*, respectively (Figure 1O). Remarkably, we found that Mut3 affects the ZNF143 binding only if it contains ZF1-2, suggesting that ZF1-2 recognizes Module4 (Figure 1P). By contrast, Mut4 appears to affect the ZNF143 binding only if it contains ZF7, suggesting that ZF7 recognizes Module1 (Figure 1Q). These data suggest a directionality of ZNF143 binding to the tandem SBS elements whereby ZF7 recognizes Module1 and ZF1-2 recognizes Module4.

### Directional ZNF143 recognition of genome-wide SBS elements

To further investigate the genome-wide directionality of ZNF143 binding without the interference of CTCF, we performed CTCF ChIP-nexus experiments (Figure 1A) and analyzed genome-wide CTCF and ZNF143 colocalization (Figure S1W). We found that ∼3/4 of ZNF143 peaks (27,150 out of 37,139) are overlapped with CBS elements (Figure 1R). Further sequence analyses of ZNF143 peaks not overlapped with CBS revealed a type of SBS elements with Module1 but lacking Module3-4 (Figure S1X)^29,34^. We investigated their recognition directionality and found that ZF7 deletion nearly abolishes the binding of ZNF143 (Figures 1S and 1T), suggesting that ZF7 plays a major role in the ZNF143 recognition of Module1. By contrast, N-terminal deletions of the first two ZFs have no effect on the ZNF143 binding (Figure 1U). Finally, mutations of single nucleotides within Module1 weaken the ZNF143 binding only if it contains ZF7 (Figures 1S and 1V). This demonstrates that ZF7 recognizes Module1. Taken together, these data suggest that ZNF143 directionally recognizes SBS elements in a flexible manner with an anti-parallel orientation (Figure 1W).

### CTCF recruits ZNF143 to *PCDH* CBS elements *in vivo*

In addition to ZNF143 enrichments at the two SBS elements of *HS5-1b1* and *HS5-1b2* within the *PCDH HS5-1* enhancer, ZNF143 is also enriched at the *PCDH* variable regions as well as super-enhancers (Figure 1A). Interestingly, ZNF143 and CTCF are colocalized at the CSE and eCBS elements within the *PCDH* variable regions as well as CBS elements within super-enhancers (Figure S2A). However, we could not find SBS motifs around these CBS elements. To see whether ZNF143 directly binds to these CBS elements, we performed comprehensive EMSA experiments using a repertoire of CSE and eCBS probes and found that ZNF143 does not bind these probes directly *in vitro* (Figure S2A). This suggests that CTCF may recruit ZNF143 to these CBS elements *in vivo*. To this end, we employed an auxin-inducible degron (AID) system^38^ to degrade CTCF *in vivo*. We first generated single-cell clones stably expressing the auxin receptor by targeting the rice *OsTIR1* gene to the human *AAVS1* (Adeno-Associated Virus Integration Site 1) locus. We then tagged CTCF alleles with AID in these cells to produce CTCF-AID clones.

We confirmed by Western blot that CTCF was degraded after 24 h of auxin treatment in CTCF-AID cells (Figure S2B). Upon CTCF degradation, there was a significant decrease of ZNF143 enrichments at the CBS elements within the variable and super-enhancer regions of the *PCDH* clusters (Figures 2A-2C). However, as internal controls, ZNF143 enrichments at the two SBS elements of *HS5-1b1* and *HS5-1b2* appear unchanged despite CTCF degradation (Figure 2C). This demonstrates that ZNF143 enrichments at the CBS elements within the *PCDH* clusters are CTCF-dependent.

**Figure 2.**
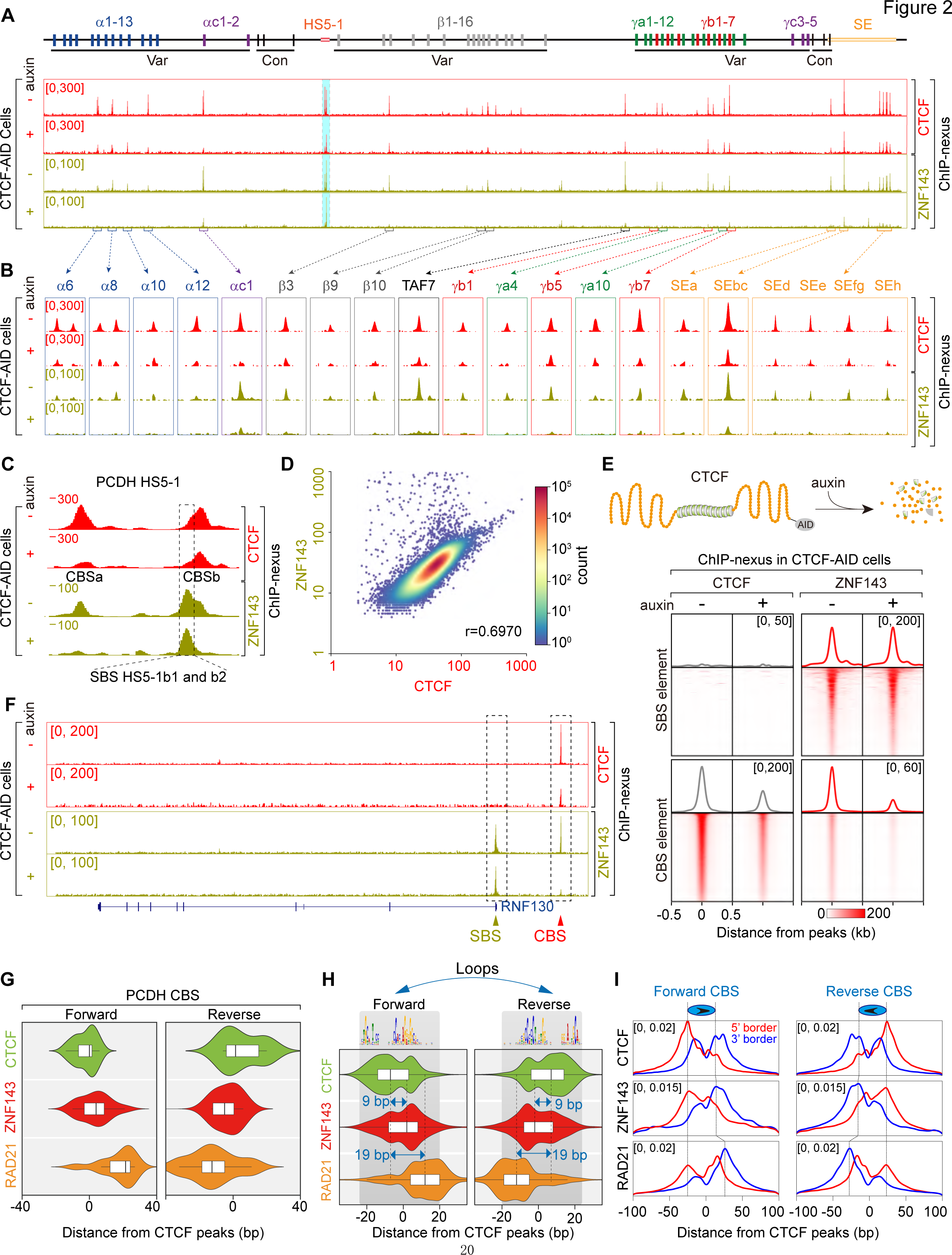
CTCF recruiting ZNF143 to CBS. (**A** and **B**) ZNF143 and CTCF ChIP-nexus profiles of the three human *PCDH* clusters in CTCF-AID cells treated with or without auxin, showing decreased ZNF143 enrichment at the *PCDH* CBS elements upon auxin-induced CTCF degradation. (C) Close-up of ZNF143 and CTCF ChIP-nexus profiles of the *HS5-1* enhancer region, showing decreased binding of ZNF143 at the *HS5-1a* and *HS5-1b* CBS elements upon auxin-induced CTCF degradation. However, there appears no decrease of ZNF143 binding at the two SBS elements of *HS5-1b1* and *HS5- 1b2*. (D) Scatter plot comparing ChIP-nexus signals of ZNF143 and CTCF at ZNF143/CTCF co-localized peaks. The correlation between ZNF143 and CTCF data is calculated using Pearson’s R. (E) Heatmaps of the CTCF and ZNF143 ChIP-nexus signals at SBS and CBS elements, respectively, with or without auxin treatment, showing decreased ZNF143 enrichments at colocalized CBS elements genome-wide upon CTCF degradation. (F) ZNF143 and CTCF ChIP-nexus profiles of the human *RNF130* locus in CTCF-AID cells treated with or without auxin, showing decreased enrichment of ZNF143 at colocalized CBS element, but not SBS, upon CTCF degradation. (**G** and **H**) Violin plots of the distributions of ZNF143, CTCF, and RAD21 ChIP- nexus peak summits of the forward or reverse CBS motifs at the *PCDH* locus (G) or genome-wide loop anchors (**H**), showing localization of ZNF143 between CTCF and RAD21 at loop anchors. (**I**) ChIP-nexus footprint profiles of ZNF143, CTCF, and RAD21 at the forward or reverse CBS elements, showing that the 3’ binding boundary of ZNF143 is located between CTCF and RAD21. Arrows indicate CBS orientations. Red or blue lines indicate 5’ or 3’ borders of footprints.

### CTCF recruits ZNF143 to CTCF sites genome-wide

We then analyzed global ZNF143 and CTCF enrichments at their colocalized CBS elements and found that ZNF143 enrichment is strongly correlated with CTCF (Figure 2D). We also found that, upon CTCF degradation, there is a significant decrease of ZNF143 enrichments at the colocalized CBS elements genome-wide (Figures 2E, 2F, and S2C). In contrast, there is no decrease of ZNF143 enrichments at the SBS elements (Figures 2E, 2F, and S2C). In conjunction with the recent finding that CTCF interacts with ZNF143 directly^36^, these data suggest that CTCF recruits ZNF143 to their colocalized CBS elements.

### ZNF143 is located between CTCF and cohesin at their colocalized loop anchors

CTCF/cohesin-mediated long-distance chromatin interactions between convergent CBS elements determine the *PCDH* promoter choice^7^. Since ZNF143 and CTCF proteins are colocalized at these forward and reverse CBS elements, we performed ChIP-nexus experiments with an antibody specifically against RAD21, a subunit of cohesin, and analyzed the colocalization of RAD21 with CTCF and ZNF143 at the *PCDH* CBS elements at single-base resolution. Interestingly, we found that ZNF143 is localized between CTCF and cohesin at both the forward and reverse *PCDH* CBS elements (Figure 2G). We then performed CTCF and RAD21 HiChIP and analyzed the global triple colocalizations of ZNF143, CTCF, and RAD21 at chromatin loop anchors. We found that ZNF143 is localized between CTCF and cohesin at loop anchors globally (Figure 2H). Further analyses of different types of CBS elements^39^ revealed that ZNF143 is localized between CTCF and cohesin for all three types of CBS elements at CTCF/cohesin loop anchors genome-wide (Figure S2D).

Previous studies revealed that RAD21 is located at ∼40 bp downstream of CTCF at loop anchors^40^. To obtain the precise localization of ZNF143 in relation to CTCF and RAD21, we analyzed DNA footprints of ZNF143, CTCF, and RAD21 at single-base resolution on their co-occupied CBS elements. We found that the 5’ borders of the three proteins are the same; however, the 3’ border of ZNF143 is slightly more inner than the 3’ border of CTCF, and the 3’ border of RAD21 is obviously localized at the innermost position (Figure 2I). In addition, loop anchors with different types of CBS elements all display the similar patterns of ZNF143, CTCF, and RAD21 footprints (Figure S2E). These data demonstrate that ZNF143 is localized between CTCF and cohesin at loop anchors.

### ZNF143 deletion compromises CTCF/cohesin-mediated chromatin looping

We next generated ZNF143 knockout single-cell clones (ΔZNF143) using CRISPR DNA fragment editing with Cas9 programmed by dual sgRNAs^10^. Western blot and RNA-seq experiments confirmed the absence of ZNF143 protein and mRNA in ΔZNF143 cells (Figures S3A and S3B). ZNF143 deletion does not affect the CTCF enrichments in the *PCDH* clusters (Figures 3A-3C).

**Figure 3.**
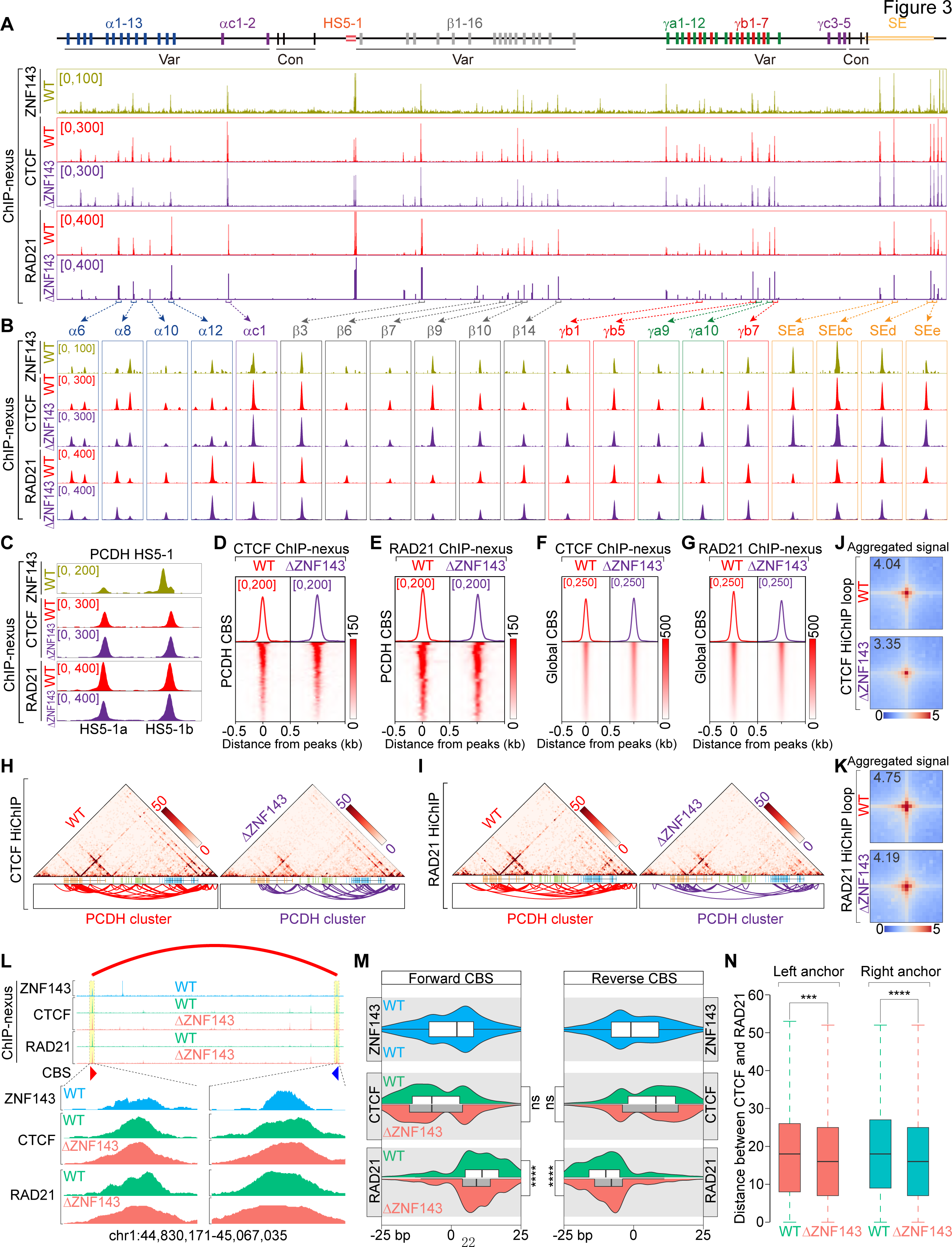
ZNF143 deletion compromising RAD21 but not CTCF enrichments. (**A**-**C**) ZNF143, CTCF, and RAD21 ChIP-nexus profiles of the *PCDH* clusters in the wild-type (WT) and ZNF143-knockout (DZNF143) cells, showing decreased enrichments of RAD21, but not CTCF, at the *PCDH* CBS elements. (**D** and **E**) Heatmaps of the CTCF (**D**) and RAD21 (**E**) ChIP-nexus signals at the *PCDH* CBS elements in WT and DZNF143 cells, showing decreased enrichments of RAD21, but not CTCF, at the *PCDH* CBS elements. (**F** and **G**) Heatmaps of the CTCF (**F**) or RAD21 (**G**) ChIP-nexus signals at genome-wide CBS elements, showing a global decrease of the enrichment levels of RAD21 (**F**), but not CTCF (**G**), at their colocalized CBS elements upon ZNF143 deletion. (**H** and **I**) CTCF (**H**) or RAD21 (**I**) HiChIP contact maps (top, at 10 kb resolution) and loops (bottom) of the *PCDH* clusters in DZNF143 cells compared to WT controls, showing decreased chromatin interactions upon ZNF143 deletion. (**J** and **K**) Aggregate peak analysis (APA) of CTCF (**J**) or RAD21 (**K**) HiChIP loops shared in both WT and DZNF143 cells, showing a decrease in loop strength upon ZNF143 deletion. (L) A shift of the RAD21 ChIP-nexus peaks toward CTCF at loop anchors upon ZNF143 deletion. Red or blue arrow indicates forward or reverse CBS motif. (M) Violin plots of the distributions of ZNF143, CTCF, and RAD21 ChIP-nexus summits at the forward or reverse CBS motifs genome-wide in WT and DZNF143 cells, showing closer proximity between RAD21 and CTCF upon ZNF143 deletion. (N) Boxplots showing deceased distances between RAD21 and CTCF peak summits upon ZNF143 deletion.

However, the RAD21 enrichments appear to decrease slightly in the *PCDH* clusters upon ZNF143 deletion (Figures 3A-3C). We then quantified CTCF and cohesin enrichments in the *PCDH* clusters using DeepTools^41^ and found a significant decrease of enrichment levels of RAD21 (p=0.006262), but not CTCF (p=0.3125), at the CBS elements within the *PCDH* clusters (Figures 3D and 3E). We then quantified global CTCF and RAD21 enrichments at their colocalized CBS elements and found a significant decrease of RAD21 (p < 2.2 × 10e-16), but not CTCF (p = 0.02143), upon ZNF143 deletion (Figures 3F and 3G).

We next performed HiChIP experiments with a specific antibody against CTCF or RAD21 and found a significant decrease of long-distance PCDH chromatin interactions upon ZNF143 deletion (Figures 3H and 3I). We then analyzed global chromatin interactions by aggregated peak analyses (APA) and found that the strengths of both CTCF and cohesin loops are weakened upon ZNF143 deletion (Figures 3J and 3K). Finally, ZNF143 deletion results in a significant decrease of chromatin loop numbers (Figure S3C), consistent with recent ZNF143 studies^36^.

### ZNF143 functions as a buffer sponge between CTCF and cohesin

We then asked whether ZNF143 deletion affects the relative locations of CTCF and cohesin at their triple colocalized sites (Figure S3D). Careful analysis of the CTCF and RAD21 ChIP-nexus data in ΔZNF143 clones revealed that the RAD21 ChIP-nexus peaks appear to be shifted toward CTCF peaks (Figures 3L and S3E). Further whole genome analyses revealed remarkably closer proximity between CTCF and RAD21 at triple colocalized CBS elements upon ZNF143 deletion (Figure 3M). In addition, this is true for all three types of CBS elements (Figure S3F). Finally, the distance between CTCF and RAD21 peaks at both left and right loop anchors of CTCF/cohesin-mediated loops also decreased upon ZNF143 deletion (Figure 3N). Together, these data suggest that ZNF143 may act as a buffer sponge between CTCF and cohesin.

### ZNF143 is required for SBS loop formation

To investigate the role of ZNF143 in loop formation, we first performed ZNF143 HiChIP experiments and predicted significant chromatin interactions with hichipper^42^ (Figure 4A). In conjunction with ZNF143 and CTCF ChIP-nexus data, we identified 2,193 SBS-SBS loops (SBS loops: both anchors at SBS elements, Table S1), 4,600 SBS-CBS loops, and 2,199 CBS-CBS loops (CBS loops: both anchors at CBS elements) identified by HiChIP experiments with a specific antibody against ZNF143 (Figures 4A, S4A, and S4B). In addition, SBS elements are preferentially located at promoter regions (Figure S4C)^29^, with over 80% of SBS promoters being active and associated with CpG islands (Figures S4D and S4E). Moreover, ∼90% of SBS loops have at least one anchor at gene promoters (Figures 4B and S4F). Furthermore, 88.7% of SBS loop-anchored promoters are active and 11.3% are inactive (Figure 4C). Finally, for SBS loops with both anchors at promoter regions, both convergent and divergent promoters could be in close spatial contacts via SBS loops (Figures 4D and 4E).

**Figure 4.**
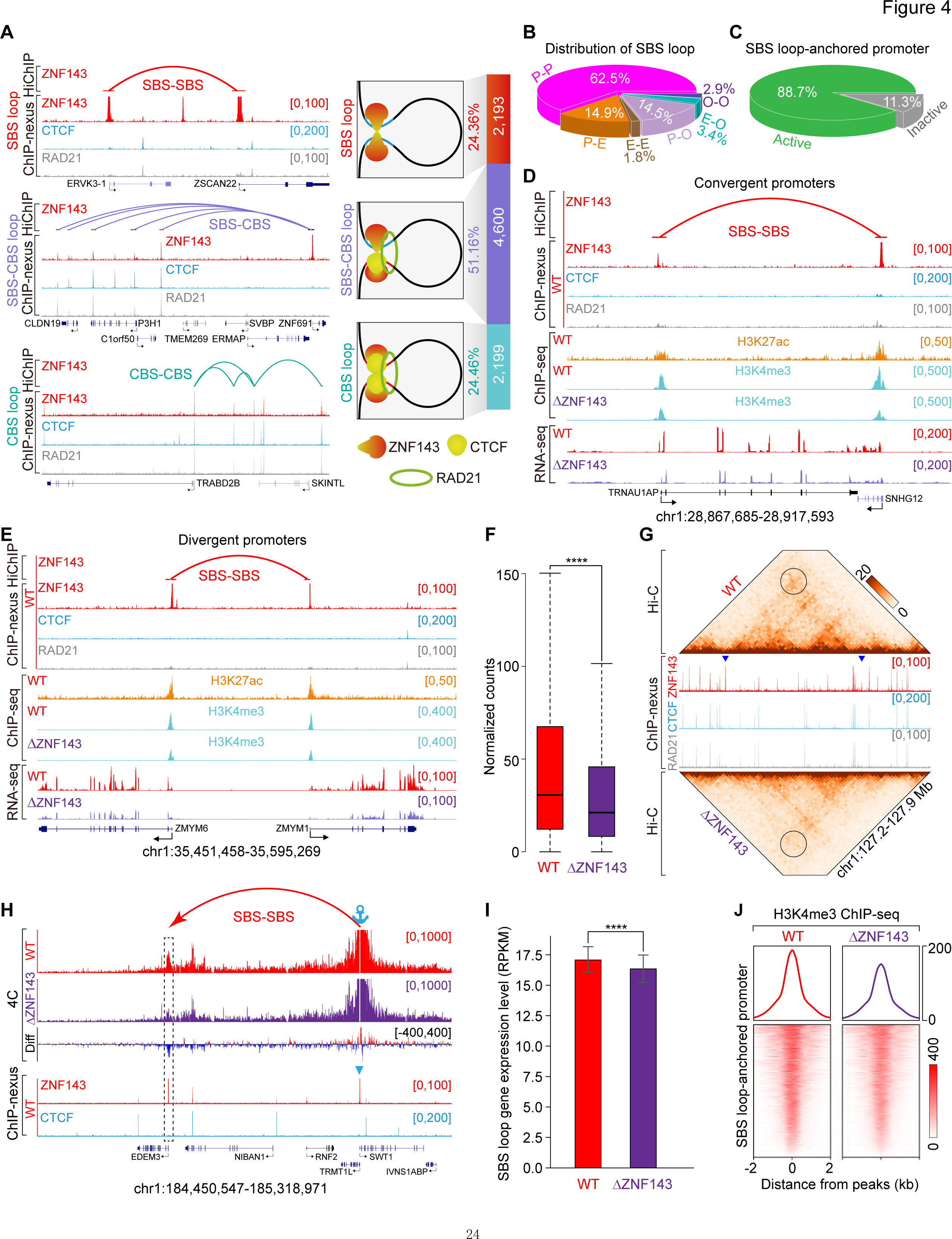
ZNF143 is required for SBS-SBS loop formation. (A) Schematics of three types of ZNF143 HiChIP chromatin loops shown by Bézier curves, including SBS loops, SBS-CBS loops, and CBS loops. (B) Distribution of SBS loops with different combinations of *cis*-regulatory elements at both anchors, showing that 91.9% of SBS loops are anchored at promoters at least on one side. (C) RNA-seq showing that the vast majority of SBS loop-anchored promoters are active. (**D** and **E**) Examples of SBS loops anchored at convergent (**D**) or divergent (**E**) promoters. (F) Normalized counts of Hi-C valid pairs of reads at the SBS loop anchors, showing a decrease of Hi-C contact strength at SBS loops upon ZNF143 deletion. (G) Hi-C contact maps centered at an SBS loop, showing decreased interactions between two SBS loop anchors upon ZNF143 deletion. Bin size, 10 kb. (H) 4C profiles confirming a significant decrease of SBS loop strength upon ZNF143 deletion. Differences are shown under the 4C profiles. (I) Decreased transcriptional levels of SBS loop-anchored genes genome wide upon ZNF143 deletion. (J) Decreased enrichments of H3K4me3 at genome wide SBS loop-anchored promoters upon ZNF143 deletion.

To further investigate the role of ZNF143 in SBS loops, we performed *in-situ* Hi-C experiments using our ΔZNF143 cell clones, with wild-type (WT) cells as controls. We analyzed valid pairs of Hi-C reads at SBS loop anchors identified by ZNF143 HiChIP experiments and found a significant decrease of loop strength upon ZNF143 deletion (Figures 4F, 4G, and S4G). Finally, to confirm the role of ZNF143 in SBS looping, we performed high-resolution 4C experiments using an SBS element as the anchor and observed a significant decrease of SBS looping upon ZNF143 deletion (Figure 4H).

To investigate the functional consequences of ZNF143 deletion, we performed RNA-seq experiments using ZNF143-deletion clones. Although there are both increased and decreased levels of gene expression upon ZNF143 deletion (Figure S4H), there is a significant decrease of expression levels of genome wide SBS loop-anchored promoters (Figure 4I). We then performed H3K4me3 ChIP-seq experiments and found a consistent decrease of H3K4me3 marks in SBS-anchored promoters (Figures 4D, 4E, 4J, and S4I). All in all, these data suggest that ZNF143 binds to promoter regions to regulate gene expression via chromatin looping.

### Aberrant large-sized SBS loops upon acute CTCF degradation

We recently found that CTCF insulators can block enhancer-promoter (E-P) contacts regardless of whether enhancers and promoters are associated with CBS elements or not^16^. To investigate whether CTCF plays a role in the specificity of ZNF143-mediated E-P or P-P chromatin looping, we performed ZNF143 HiChIP experiments in auxin-treated and untreated CTCF-AID cells. As expected, CBS loop number is substantially decreased upon addition of auxin (Figure S5A)^43–45^. The CBS loops identified by ZNF143 HiChIP still obey the forward-reverse convergent rule (Figure S5B). However, the size of SBS loops appears larger in auxin-treated than -untreated cells (Figures 5A and S5C). In addition, acute CTCF degradation leads to a global decrease of chromatin CBS loop numbers (Figure S5D). Interestingly, both the number and average size of SBS loops are increased (Figures 5B and 5C), while the average size of CBS loops identified with ZNF143 HiChIP does not change (Figure S5E). Remarkably, there is a strong shift from small SBS loops to large SBS loops upon CTCF degradation (Figures 5D-5F and S5F), suggesting that CTCF insulators have a role in maintaining or restraining proper SBS loop size.

**Figure 5.**
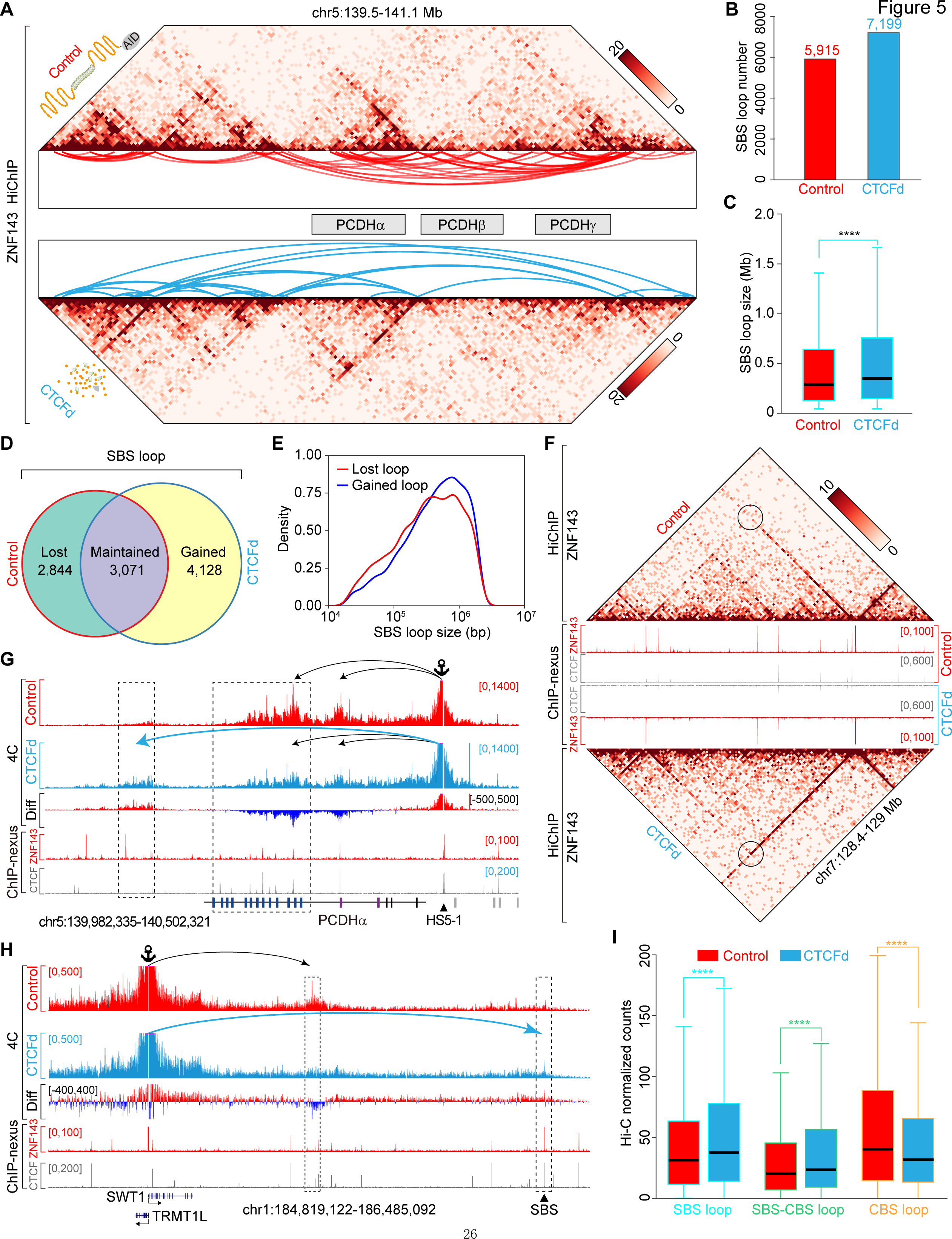
Formation of aberrant large-sized SBS loops upon CTCF degradation. (A) ZNF143 HiChIP contact map at the *PCDH* clusters in auxin-treated (CTCF degraded, CTCFd) or -untreated (control) CTCF-AID cells, showing the formation of aberrant large-sized loops upon CTCF degradation. Bin size, 10 kb. (B) SBS loop number in CTCFd cells compared to controls, showing an increase of SBS loops upon CTCF degradation. (C) Increase of SBS loop sizes upon CTCF degradation. (D) Number of SBS loops that are lost, maintained, or gained after CTCF degradation. (E) Size distribution of SBS loops that are lost or gained upon CTCF degradation, showing increased size of gained SBS loops. (F) ZNF143 HiChIP heatmaps showing increased interactions between SBS loop anchors upon CTCF degradation. Bin size, 10 kb. (**G** and **H**) 4C profiles confirming a significant decrease of small loop strengths and a significant increase of aberrant large loops upon CTCF degradation. Differences are shown under the 4C profiles. (I) Normalized counts of Hi-C valid pairs of reads for three types of ZNF143 HiChIP loops, showing a significant increase of SBS-anchored loops and decrease of CBS loops upon CTCF degradation.

To confirm this observation, we performed 4C at the *PCDHα* and *SWT1/TRMT1L* loci and found that CTCF depletion results in a significant decrease of small loops but a significant increase of large loops (Figures 5G and 5H). Finally, we analyzed valid pairs of Hi-C reads for both CTCF-depleted and un-depleted cells across all three types of ZNF143-HiChIP loops and found a significant increase of both SBS and SBS-CBS loop strengths, but a significant decrease of CBS loop strength, upon CTCF degradation (Figure 5I). Overall, these data suggest that CTCF may function as an insulator to prevent the aberrant formation of large-sized SBS loops.

### ZNF143 deletion alters TADs and compartments

To investigate the role of ZNF143 in higher-order chromatin organization, we performed *in-situ* Hi-C experiments in our ΔZNF143 single-cell clones. We observed a significant decrease of chromatin contacts on chromosome 5 and the *PCDH* clusters upon ZNF143 deletion (Figures 6A-6C and S6A). Virtual 4C with *HS5-1* as the anchor showed reduced chromatin interactions with *PCDHα* alternate variable exons (Figure 6D). We then deleted the two tandem SBS elements within the *HS5-1* enhancer (Figure 6E) and performed 4C experiments with *HS5-1* as the anchor. We observed a similar decrease of chromatin interactions with *PCDHα* alternate variable exons (Figure 6F).

**Figure 6.**
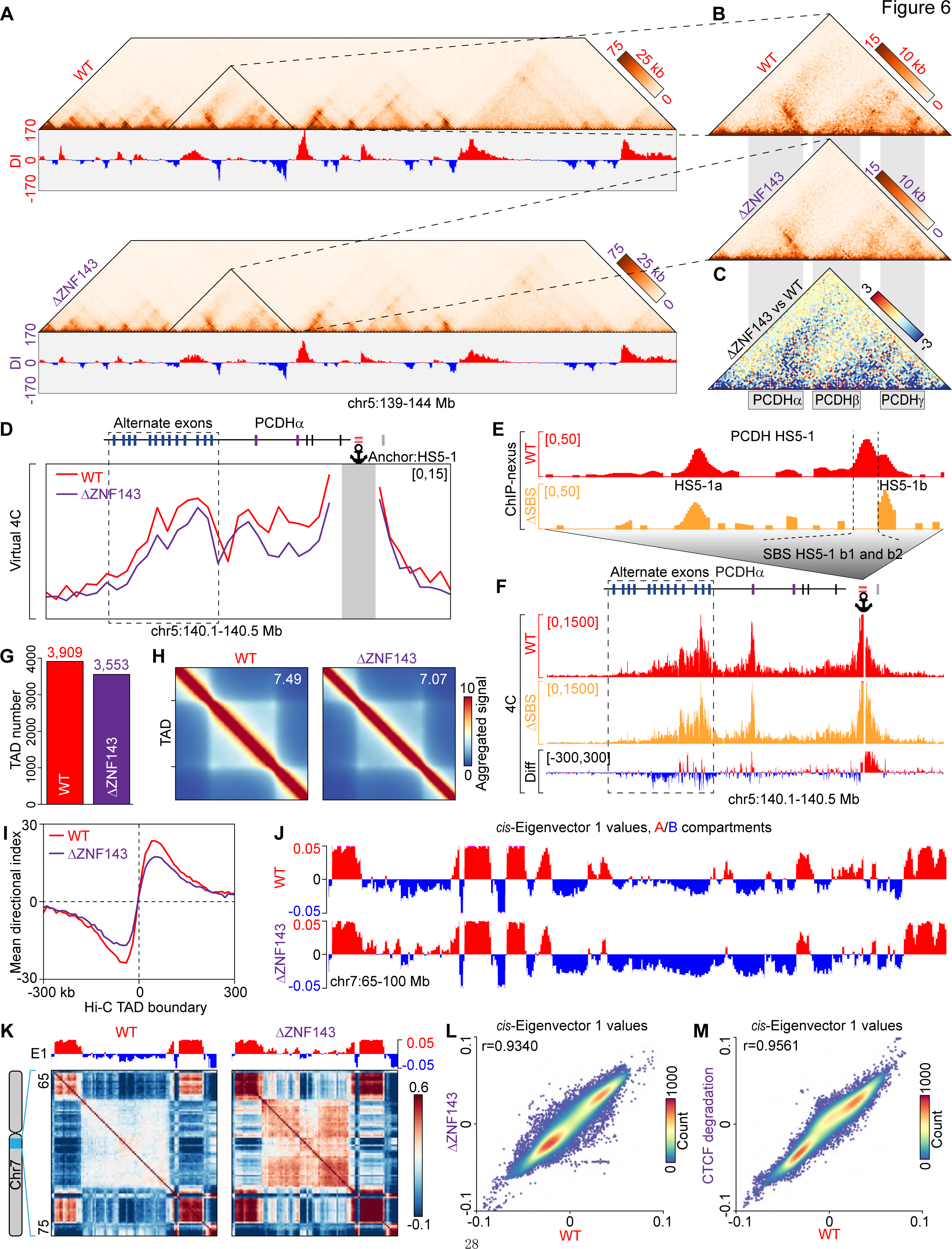
ZNF143 deletion weakens TAD boundary. (A) Hi-C contact maps of the chr5:139-144 Mb region at 25 kb resolution with directionality index (DI). (B) Close-up of the *PCDH* clusters at 10 kb resolution. (C) The differential contact map between DZNF143 and WT cells at the *PCDH* clusters, showing a significant decrease of chromatin contacts upon ZNF143 deletion. (D) Virtual 4C profiles using the *HS5-1* enhancer as an anchor, showing a decrease of chromatin interactions with the *PCDHa* alternative exons upon ZNF143 deletion. (E) ZNF143 ChIP-nexus profiles of the *HS5-1* enhancer, showing disappeared signals at the deleted regions of the two tandem SBS elements but not at the flanking CBS elements in SBS-deleted (DSBS) cell clones. (F) 4C profiles using *HS5-1* as an anchor showing decreased interactions with the *PCDHa* alternative exons with SBS deletion. Interaction differences are shown under the 4C profiles. (G) TAD number in WT or DZNF143 cells. (H) Aggregate domain analysis (ADA) of TADs showing a decrease of intra-TAD contacts upon ZNF143 deletion. (I) Profiles of the average DI values in 300 kb regions flanking all TAD boundaries in DZNF143 cells compared to WT cells, showing weakened TAD insulation upon ZNF143 deletion. (J) *cis-*Eigenvector 1 values at chr7:65-100 Mb in WT or DZNF143 cells. The regions with value above zero are compartment A, and those with value below zero are compartment B. (K) *cis-*Eigenvector 1 (E1) values and heatmaps of Pearson’s correlation at chr7:65-75 Mb, showing a finer compartmentalization upon ZNF143 deletion. (**L** and **M**) Scatter plot showing the Pearson’s correlation of *cis-*Eigenvector 1 values between WT and DZNF143 (**L**), or between WT and CTCFd (**M**).

Global analyses of Hi-C data from ΔZNF143 cell clones revealed decreased TAD number (Figure 6G), intra-TAD contacts (Figure 6H), and loop strength (Figure S6B). In addition, ZNF143 deletion significantly weakens the strength of TAD boundaries (Figures 6I, S6C, and S6D) probably because CTCF/ZNF143 co-occupied CBS elements tend to be located at TAD boundaries (Figure S6E). These data suggest that ZNF143 plays an important role in TAD boundary formation.

We next investigated the effect of ZNF143 deletion on chromatin segregation between compartments A and B. Compartment signals (Figure 6J) and Pearson’s correlation heatmaps (Figure 6K) suggest that ZNF143 deletion leads to transitions from A to B compartments or vice versa. Further correlation analysis revealed that contact maps (Figure S6F) and compartments signals are altered upon ZNF143 deletion (Figure 6L) compared to almost no alteration upon CTCF degradation (Figure 6M). In summary, our genetic experiments demonstrate that ZNF143 plays an important role in higher-order chromatin organization.

### ZNF143 and CTCF collaborate to organize 3D genome architecture

To systematically investigate the integrated role of ZNF143 and CTCF in 3D genome organization, we screened for ZNF143-deleted single-cell clones in CTCF-AID cells using CRISPR DNA-fragment editing (Figure 7A). We then performed *in-situ* Hi-C using ZNF143-deleted and CTCF-depleted (abbreviated as CTCFd) cells by treatment with auxin (Figures 7A and S7A). We found that deletion of ZNF143 in CTCF-depleted cells results in a further decrease in TAD number (Figure S7B), intra-TAD contacts in PCDH clusters and across the entire genome (Figures 7B, and 7C), loop strength (Figure 7D), and boundary strength (Figures 7E and S7C) compared to CTCF depletion alone. This suggests that ZNF143 and CTCF collaborate in forming chromatin loops.

**Figure 7.**
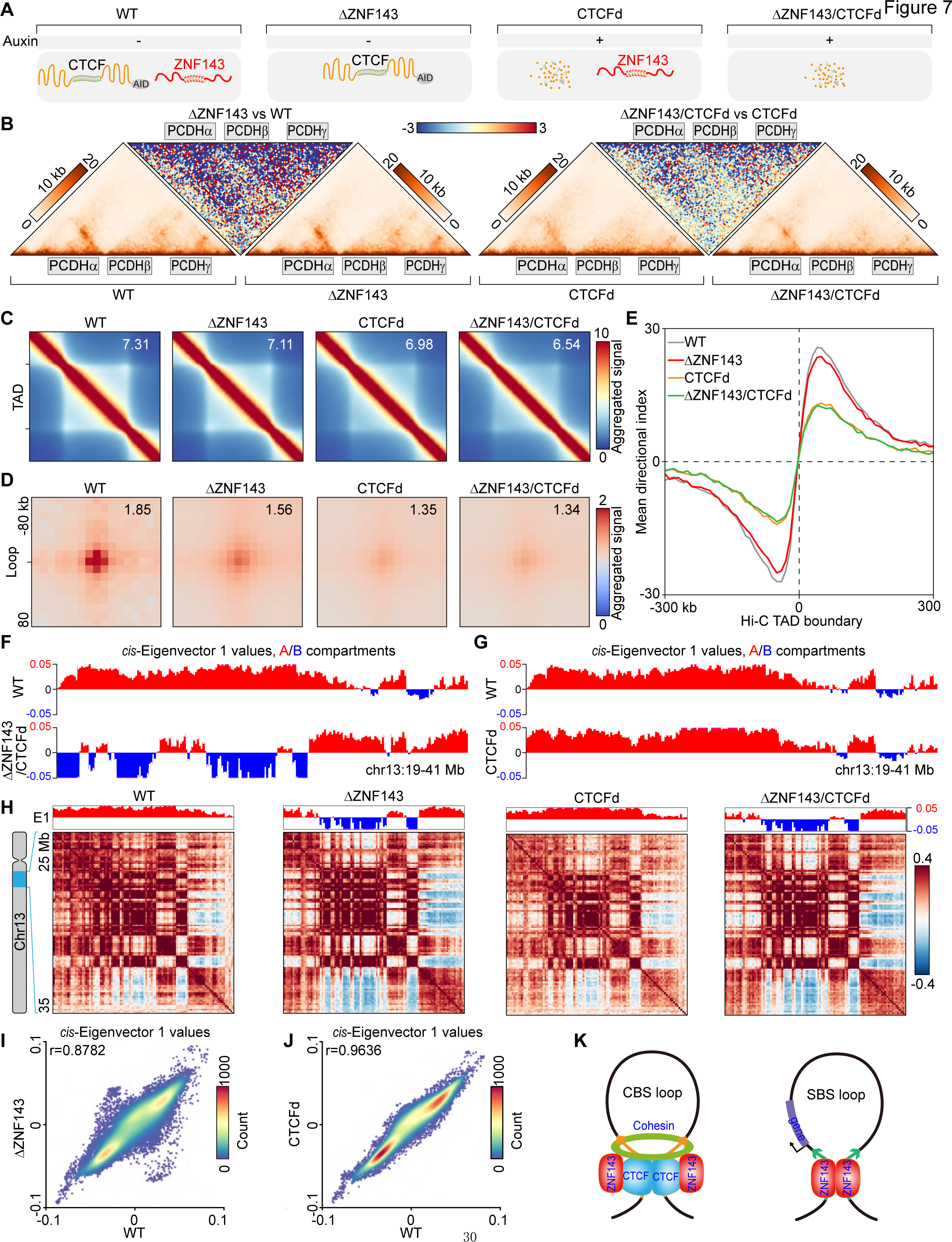
ZNF143 and CTCF collaborate to organize 3D genome architecture. (A) Schematics of four distinct cell populations used for *in-situ* Hi-C, CTCF-AID cells treated with (CTCFd) or without (WT) auxin, ZNF143-deleted CTCF-AID cells treated with (DZNF143/CTCFd) or without (DZNF143) auxin. (B) Hi-C contact maps at the *PCDH* clusters showing a decrease of intra-TAD contacts by not only CTCF degradation but also ZNF143 deletion. Differential heatmap is shown between DZNF143 and WT, or between DZNF143/CTCFd and CTCFd. (**C** and **D**) ADA plot for all TADs (**C**) or APA plot for all loops (**D**) called from Hi- C data of WT, DZNF143, CTCFd, or DZNF143/CTCFd cells, showing a global decrease in intra-TAD contacts or loop strengths upon both CTCF degradation and ZNF143 deletion. The ADA or APA score is indicated in the upper right corner of each panel. (**E**) Profiles of the average DI values in 300-kb regions flanking all TAD boundaries in WT, DZNF143, CTCFd, or DZNF143/CTCFd cells, showing weakened TAD insulation upon both CTCF degradation and ZNF143 deletion. (**F** and **G**) *cis-*Eigenvector 1 values at chr13:19-41 Mb in DZNF143/CTCFd compared to WT cells (**F**), or in CTCFd compared to WT cells (**G**), showing an alteration in compartmentalization upon ZNF143 deletion but not CTCF degradation. (H) *cis*-Eigenvector 1 (E1) values and heatmaps of Pearson’s correlation at chr13:25-35 Mb, showing a finer compartmentalization upon ZNF143 deletion but not CTCF degradation. (**I** and **J**) Scatter plot of the Pearson’s correlation of *cis-*Eigenvector 1 values showing decreased correlation upon ZNF143 deletion (**I**) but not CTCF degradation (**J**). (**K**) Model of dichotomic ZNF143 function in chromatin looping.

We observed a remarkable alteration in chromatin compartmentalization in ZNF143-deleted and CTCF-depleted cells (Figure 7F), in striking contrast to no compartmentalization alteration with acute CTCF depletion (Figure 7G)^43,44^. Pearson’s correlation map revealed a similar alteration of compartmentalization upon ZNF143 deletion but not acute CTCF depletion (Figure 7H). Consistently, contact maps and compartment signals are altered upon ZNF143 deletion (Figures 7I, S7D, and S7E) but not acute CTCF depletion (Figures 7J, S7D, and S7E). These data revealed prominent distinctions between ZNF143 and CTCF in genome compartmentalization.

## Discussion

CTCF/cohesin-mediated directional chromatin contacts between remote super- enhancers and target promoters via continuous active “loop extrusion” determine promoter choice of clustered *Pcdh* genes^7,16,25^. Developmental regulation of their higher-order chromatin organization in distinct cell-types in the brain enables neurons to achieve proper serotonergic axonal tiling and olfactory sensory neuronal axon convergence, as well as neocortical fine spatial arrangement and connectivity^46,47^. Using the *PCDH* clusters as model genes, we uncovered that ZNF143 mediates chromatin interactions in both CTCF- dependent and -independent manners. We first mapped the precise locations of SBS elements within the *PCDHa HS5-1* enhancer. By comprehensive gel shift experiments, we found that the central zinc-finger domain of ZNF143 directionally recognizes the *HS5-1* tandem SBS elements in an antiparallel manner. In addition, although CTCF directly interacts with the CES (Conserved essential surface) of cohesin^13–15^, our data suggest that, at the triple ZNF143, CTCF, and cohesin colocalized CBS elements, the ∼30 CTCF residues between its cohesin-CES-binding segment and DNA-binding domain are flexible and that ZNF143 is located between CTCF and cohesin as a buffer sponge because genetic deletion of ZNF143 results in narrowing of the space between them. This is consistent with the flexible linker model for cohesin engaging the N-terminus of CTCF and their relative mapping positions^13,40^. Finally, acute CTCF degradation leads to a decrease of ZNF143 enrichment, while ZNF143 deletion does not alter CTCF enrichment but slightly reduces cohesin. Thus, ZNF143 may function to keep CTCF and cohesin appropriately spaced in 3D geometry and to stabilize cohesin anchoring at CBS elements.

Interphase genomes are organized into complex higher-order structures including chromatin loop, TAD and subTAD, and chromatin compartment. Among numerous proteins involved in higher-order genome organization, ZNF143 is interesting in that it is largely colocalized with CTCF and is also independently located in gene promoters^29,48,49^. Hi-C experiments with ZNF143-deleted CRISPR single-cell clones demonstrated that ZNF143 is required for the formation of SBS loops. In addition, ZNF143 HiChIP experiments, in conjunction with acute CTCF degradation, revealed that CTCF depletion results in the formation of aberrantly-large-sized SBS loops, suggesting that CTCF insulators play a role in maintaining proper sizes of SBS loops. Moreover, genetic deletion of ZNF143, in conjunction with *in situ* Hi-C experiments, showed that ZNF143 is crucial for chromatin compartmentalization. Previous studies suggested that ZNF143 is mainly associated with E2F-bound promoters and CTCF-bound enhancers^28,29,33,34,36,37,48–51^. ZNF143 might regulate genome domains of A/B compartments by altering activity of SBS-associated promoters. Finally, we found that ZNF143 and CTCF collaborate to regulate higher-order chromatin organization.

Recent studies suggest that ZNF143 mediates the formation of short-range chromatin loops^36,37^; however, its DNA recognition mechanism and role in loop formation remain unclear. Although ZNF143 and CTCF colocalize at CBS elements, our EMSA experiments suggest that ZNF143 does not directly bind CBS elements. In addition, our ChIP-nexus data showed that CTCF depletion decreases ZNF143 enrichments at CBS elements, suggesting that CTCF recruits ZNF143 to CBS elements. We propose a looping model for dichotomic ZNF143 functions in genome architecture (Figure 7K). At CBS elements, ZNF143 stabilizes CTCF-cohesin interactions to regulate chromatin interactions via cohesin “loop extrusion”. ZNF143 also directly binds SBS elements in an antiparallel manner to regulate promoter-promoter contacts. Through activating SBS-associated promoters, ZNF143 can influence higher- order A/B compartmentalization.

## Acknowledgments

This work was supported by grants from the National Natural Science Foundation of China (32330016), the National Key R&D Program of China (2022YFC3400200) and the Science and Technology Commission of Shanghai Municipality (21DZ2210200).

## Author contributions

Q.W. conceived the research. M.Z. performed experiments. M.Z., J.L., and H.H. analyzed data. M.Z., H.H., and Q.W. wrote the manuscript.

## STAR★Methods

### Resource availability

#### Lead contact

Further information and requests for resources and reagents should be directed to and will be fulfilled by the lead contact, Qiang Wu.

#### Materials availability

Material generated in this study is available from the Lead contact.

### Experimental model and study participant details

#### Cell culture

Human HEC-1-B cells were cultured in MEM with L-glutamine and Earle’s balanced salts (Hyclone) supplemented with 1 mM sodium pyruvate (Sigma), 1% penicillin-streptomycin (Gibco), and 10% fetal bovine serum (Gibco). HCT116 cells containing the OsTIR1 gene integrated at the AAVS1 locus and the mAID-mClover tag integrated after the last codon of the RAD21 gene (RAD21-mAC HCT116)38 were cultured in RPMI 1640 medium with L-glutamine (Gibco) supplemented with 1% penicillin-streptomycin and 10% fetal bovine serum. All cells were maintained at 37°C in a humidified 5% CO2 incubator. For degradation of AID-tagged proteins, OsTIR1-expressed cells were treated with 500 μM indole-3-acetic acid (IAA, auxin) for 24 h.

### Method details

#### Plasmid construction

To construct the donor plasmid for OsTIR1 expression from the AAVS1 locus, a codon-optimized OsTIR1 gene was amplified from the genomic DNA of RAD21-mAC HCT116 cells by PCR and cloned into the Spe I and Bam HI sites under the control of the CMV immediate early promoter (*P*CMV-IE) in the pLVX-IRES-Puro plasmid (Clontech). The upstream (AAVS1U) and downstream (AAVS1D) homology arms were amplified from the HEC-1-B genomic DNA by PCR. AAVS1U, *P*CMV-IE-OsTIR1-IRES-Puro, and AAVS1D were then assembled and cloned into pGEM-T-Easy (Promega) to generate the pGEM-AAVS1- OsTIR1-Puro donor plasmid.

To construct the CTCF-AID donor plasmid, a 68-aa mAID tag was amplified from the RAD21-mAC HCT116 genomic DNA by PCR. Homology arms were designed to allow in-frame C-terminal fusion of mAID to the CTCF gene. The upstream homology arm (CTCF-U) corresponds to the upstream sequence of the CTCF stop codon. The downstream homology arm (CTCF-D) corresponds to the sequence downstream of the last CTCF codon. CTCF-U, mAID, and CTCF-D were assembled and cloned into pGEM-T-Easy to generate the pGEM-CTCF-AID donor plasmid.

The sgRNA expression plasmids targeting *AAVS1*, *CTCF,* or *ZNF143* were generated as previously described^52^. Briefly, target-specific sgRNA oligos were ordered and cloned into the Bsa I site of pGL3-U6-sgRNA-PGK-Puro for sgRNA transcription. All primers used are listed in Supplementary Table 2. All plasmid constructs were confirmed by Sanger sequencing.

#### Preparation of truncated zinc finger domains of ZNF143

Full-length coding sequences (CDS) of human ZNF143 were amplified with PCR from a cDNA library of human cells. Each truncated ZNF143 containing different combinations of zinc fingers (ZF) was fused with myc-tag sequence at 5’ end (ZF1-7, ZF1-6, ZF1-5, ZF1-4, ZF1-3) or at 3’ end (ZF1-7, ZF2-7, ZF3-7, ZF4-7, ZF5-7) through amplification with PCR from the full length ZNF143 CDS with specific primers (Supplementary Table 2) and cloned into the Eco RI and Not I/Xba I sites of the pTNT vector (Promega L5610) under the control of the T7 promoter. Each construct was confirmed by Sanger sequencing and used as DNA template to generate corresponding polypeptide of truncated ZNF143 *in vitro* using TNT T7 Quick Coupled Transcription/Translation Systems (Promega L1170) according to the manufacturer’s instructions. For each reaction, 200 ng of pTNT plasmid was mixed with 8 μL of TNT Quick Master Mix and 0.2 μL of 1 mM methionine and diluted with H_2_O to a final reaction volume of 10 μL. The reaction mixture was incubated at 30°C for 90 min. The synthesized protein was confirmed by the Western blot using an anti-myc-tag antibody (Millipore 05-724), aliquoted, and stored at -80°C.

#### Electrophoretic mobility shift assay (EMSA)

EMSA experiments were performed as previously described^10,69^ with slight modification. Briefly, each DNA fragment containing probe sequences was amplified from the human genomic DNA with PCR and cloned into the pGEM-T easy vector to generate the template plasmid. The resulting template plasmid was used to generate mutated template plasmid using the Q5 Site-Directed Mutagenesis Kit (NEB E0554S). All of the generated template plasmids were confirmed by Sanger sequencing. Each Probe was generated by PCR using a 5’ biotin-labeled forward primer and a reverse primer^10^ (Supplementary Table 2) and gel-purified. Probe concentration was measured with NanoDrop (Thermo) and adjusted to 100 fmol/μL.

The LightShift Chemiluminescent EMSA Kit (Thermo 20148) was then used for the EMSA experiments. The probe was incubated with 1x binding buffer, 0.05 μg/μL poly (dI-dC), 2.5% (v/v) glycerol, 0.1% NP-40, 5 mM MgCl2, 0.1 mM ZnSO4, and the synthesized protein at room temperature for 20 min.

For supershift, an anti-myc tag antibody was added and the mixture was incubated for an additional 20 min at room temperature. After incubation, the reaction mixture was separated on a 5% nondenaturing polyacrylamide gel which was pre-electrophoresed for 1 h in ice-cold 0.5x TBE buffer (45 mM Tris-borate, 1 mM EDTA, pH8.0). After transferring to a nylon membrane and crosslinking, the membrane was blocked with a 1x blocking buffer for 10 min at room temperature, washed three times with 1x washing buffer, and sequentially incubated with substrate equilibration buffer for 10 min at room temperature and Stabilized Streptavidin-Horseradish Peroxidase Conjugate. ChemiDoc XRS+ system (Bio-Rad) was used to detect the biotin-probe.

#### CRISPR screening of single-cell ZNF143-deletion clones

The ZNF143-deletion cell clones were generated using CRISPR/Cas9- mediated DNA-fragment editing^10,52^ with dual sgRNAs targeting exons 6 and 11 of the endogenous ZNF143 gene. Briefly, HEC-1-B cells were cultured in 6-well plates until ∼80% confluent. Cells were then transfected with 1 µg pcDNA3.1- Cas9 and 1 µg dual sgRNA expression plasmids per well using Lipofectamine 3000 (Invitrogen). Transfected cells were cultured in medium containing 2 μg/mL puromycin for 3 days. After puromycin selection, the cells were dissociated with trypsin, resuspended, diluted, and seeded into 96-well plates at roughly 1 cell per well. After ∼2 weeks of culturing, single cell clones were picked under a microscope and genotyped by PCR using specific primers (Supplementary Table 2). Positive clones were confirmed by Sanger sequencing. The loss of ZNF143 protein in the knockout cells was also confirmed by Western blot using an anti-ZNF143 antibody (Abcam). We obtained two homozygous ZNF143 deletion clones which grow very slow because ZNF143 is known to regulate cell cycle gene expression^49^.

#### Western blot

Cells within the 6-well plate were washed once with PBS and lysed in the RIPA buffer (50 mM Tris-HCl pH 7.4, 150 mM NaCl, 1% Triton X-100, 1% sodium deoxycholate, 0.1% SDS) containing 1x protease inhibitors. The protein lysates were then denatured at 100°C for 10 min. Protein concentration was determined by the BCA protein assay. Equal amounts of protein were separated on SDS-PAGE and transferred to a nitrocellulose membrane. After blocking with 5% fat-free dry milk in the PBST (PBS containing 0.1% Tween-20) buffer for 1 h at room temperature, the membrane was washed three times with PBST buffers and incubated with the primary antibody at 4°C overnight with slow rotation. After washed three times with the PBST buffer, the membrane was finally incubated with the Fluorescent-dye conjugated secondary antibody for 2 h at room temperature, washed four times with the PBS buffer, and scanned by the Odyssey System (LI-COR Biosciences).

#### ChIP-nexus

ChIP-nexus experiments were performed as previously described^39^ with slight modification. For each ChIP-nexus experiment, 1x 10^7^ cells were crosslinked with 1% formaldehyde and quenched with glycine. Crosslinked cells were lysed twice with 1 mL of the ChIP buffer (10 mM Tris-HCl pH 7.5, 1 mM EDTA, 1% Triton X-100, 0.1% sodium deoxycholate, 150 mM NaCl, 1x protease inhibitors) and sonicated for DNA fragmentation. After centrifugation at 12,000 rpm for 10 min, the supernatant was transferred into a new tube and incubated with specific antibody overnight at 4°C with slow rotation. Magnetic protein A/G beads were added to enrich the targeted chromatin. The beads were washed to remove the nonspecific binding. The enriched DNA was eluded and purified for library preparation.

The DNA ends were blunted using NEBNext End Repair Module (NEB E6050S), added with dATP using Klenow exo^-^ (NEB M0212S), and ligated to specific adaptors (Nex_adapter_UbamHI and Nex_adapter_BN5BamHI, Supplementary Table 2). The 5’ overhang of the adaptor-ligated DNA was filled using Klenow exo^-^ (NEB M0212S) and T4 DNA polymerase (NEB M0203S).

The generated blunt-end DNA was digested with Lambda Exonuclease (NEB M0262S) and RecJf exonuclease (NEB M0264S). After reverse-crosslinking at 65°C overnight, DNA was extracted using phenol/chloroform and precipitated with ethanol containing sodium acetate and glycogen. The purified DNA was denatured to generate ssDNA, which was self-circularized with ssDNA Ligase (Epicenter CL4111K). The circular DNA was annealed with an oligonucleotide (Nex_cut_Bam HI Supplementary Table 2) containing the Bam HI site. After digestion with Bam HI and re-precipitation with ethanol, the DNA was used for construction of the ChIP-nexus library. The DNA library was purified using gel-purification and sequenced on an Illumina platform.

#### CRISPR screening of single-cell CTCF-AID clones

The CTCF-AID HEC-1-B cells were generated in two steps using CRISPR/Cas9-mediated editing. In the first step, an OsTIR1 expression cassette (*P*_CMV-IE_-OsTIR1-IRES-Puro) was integrated into the AAVS1 safe harbor locus of the HEC-1-B cells to generate the parental cell line. In the second step, an AID cassette was fused to CTCF upstream of the stop codon in the CTCF gene of the OsTIR1-expressing parental cell.

For each editing step, cells were cultured in 6-well plates until ∼80% confluent. Cells were then transfected with 1 µg of pcDNA3.1-Cas9, 0.5 µg of the donor plasmid, and 0.5 µg of sgRNA expression plasmid per well using Lipofectamine 3000 (Invitrogen). Transfected cells were cultured in medium containing 2 μg/mL puromycin for 3 days. After puromycin selection, the cells were dissociated with trypsin, resuspended, diluted, and seeded into 96-well plates. After ∼2 weeks of continuous culturing, single-cell clones were picked under a microscope and genotyped by PCR using specific primers (Supplementary Table 2). Positive clones were confirmed by Sanger sequencing.

#### RNA-seq

For each RNA-seq experiment, about one million cells were washed twice with PBS, lysed with 1 mL Trizol (Invitrogen 15596026) for 15 min at room temperature, supplemented with 0.2 mL chloroform, vortexed, incubated for 2- 3 minutes at room temperature, and centrifuged at 12,000 rpm for 15 min at 4°C. Aqueous phase was transferred to a new tube, mixed with 0.5 mL isopropanol, incubated at room temperature for 10 min, and centrifuged at 12,000 rpm for 10 min at 4°C to obtain the total RNA pellet. To remove salts, the pellet was washed with 75% ethanol. Finally the pellet was dissolved in 100 μL of nuclease-free water. The total RNA was purified using the RNeasy kit (QIAGEN 75142) to remove residual DNA. One µg purified RNA was used to extract mRNA with NEBNext poly(A) mRNA Magnetic Isolation Module (NEB E7490S). RNA-seq library was constructed using NEBNext Ultra II RNA Library Prep Kit and sequenced on an Illumina platform.

#### ChIP-seq

For each ChIP-seq experiment, 10-20 million cells were collected and washed twice with PBS, digested with trypsin, resuspended in 10 mL medium. Formaldehyde (Thermo 28908) was added to a final concentration of 1% for cross-linking at room temperature for 10 min. The glycine was added to a final concentration of 125 mM and incubated at room temperature for 5 min to quench the cross-linking reaction. Cross-linked cells were centrifuged at 2,500 g for 10 min at 4°C. The cell pellets were washed with ice-cold PBS. Cells were lysed twice using 1 mL ice-cold ChIP buffer 1 (10 mM Tris-HCl pH 7.5, 1 mM EDTA, 1% Triton X-100, 0.1% sodium deoxycholate, 150 mM NaCl, 1x protease inhibitors) at 4°C for 10 min with slow rotation, spun at 2,500 g for 5 min at 4°C to obtain cell nuclei.

The isolated nuclei were resuspended in 0.7 mL ChIP buffer 1 followed by incubation on ice for 10 min and were sonicated in a non-contact manner with Bioruptor Plus Sonicator (Diagenode) at high intensity for 30 rounds of 30 sec on / 30 sec off to generate 100-10,000 bp DNA fragments. The sonicated samples were spun at 14,000 g for 10 min at 4°C. The supernatants were transferred to a new tube and precleared with 50 μL agarose protein A beads (16-157 Millipore). The primary antibody was added and incubated overnight at 4°C with slow rotation for immunoprecipitation. The agarose protein A beads (50 μL) were added and incubated at 4°C with slow rotation for 3 h. The samples were spun at 2,000 g for 1 min and followed by sequential washing with ChIP buffer 1, ChIP buffer 2 (10 mM Tris-HCl pH 7.5, 1 mM EDTA, 1% Triton X-100, 0.1% sodium deoxycholate, 400 mM NaCl), ChIP buffer 3 (10 mM Tris-HCl pH 7.5, 1 mM EDTA, 1% Triton X-100, 0.1% sodium deoxycholate), and ChIP buffer 4 (50 mM HEPES pH 7.5, 1 mM EDTA, 1% NP-40, 0.7% sodium deoxycholate, 500 mM LiCl). The washed antibody/protein/DNA complexes were eluted twice with 100 μL elution buffer (50 mM Tris-HCl pH 8.0, 10 mM EDTA, 1% SDS) by incubation at 65°C for 30 min with vortexes. The 200 μL eluted solutions were mixed with 200 μL TE buffer, de-cross-linked at 65°C overnight with vortexes, and sequentially digested with 2 μL RNase A at 37°C for 2 h and with 8 μL proteinase K at 55°C for 2 h. The DNA was purified with 400 μL phenol/chloroform, precipitated, and resuspended in 20 μL nuclease-free water. DNA concentration was measured by PicoGreen reagents. 10 ng DNA was used for library construction using NEBNext Ultra II DNA Library Prep Kit for Illumina (NEB). Libraries were sequenced on an Illumina HiSeq platform.

#### HiChIP

We performed HiChIP experiments as described recently^26^. Briefly, for each HiChIP experiment, ∼10 million cells were collected, spun down, and resuspended in 10 mL fresh medium after centrifugation at 500 g. The cells were crosslinked with 1% formaldehyde for 10 min at room temperature with slow rotation and the crosslinking reaction was quenched with 125 mM glycine.

The crosslinked cells were spun down at 800 g for 5 min, washed once with ice-cold PBS, and lysed twice with 1 mL ice-cold lysis buffer (10 mM Tris-HCl pH 8.0, 10 mM NaCl, 0.2% NP-40, 1x protease inhibitors) for 15 min at 4°C with slow rotation to obtain nuclei. The samples were spun down at 2,500 g at 4°C for 5 min. The pellets were resuspended in 100 μL of 0.5% SDS solution and incubated at 62°C for 10 min. 285 μL of H2O and 50 μL of 10% Triton X-100 were added and incubated at 37°C for 15 min. 50 μL of 10 × NEBuffer 2 and 300 U of Mbo I were then added and rotated at 37°C for 2 h. To fill in the restriction fragment overhangs and label the DNA ends with biotin, 29.5 μL of mixture (15 μL of 1 mM biotin-14-dATP, 1.5 μL of 10 mM dGTP, 1.5 μL of 10 mM dCTP, 1.5 μL of 10 mM dTTP and 50 U of the Large Klenow Fragment of DNA Polymerase I) was added and rotated at 37°C for 1 h. The ligation mix (660 μL of H2O, 150 μL of 10 × T4 DNA ligase buffer, 125 μL of 10% Triton X- 100, 3 μL of 50 mg/ml BSA and 400 U of T4 DNA ligase) were then added and rotated at room temperature for 4 h. The nuclei were pelleted at 2,500 g for 5 min at room temperature and the supernatant was removed. The samples were sonicated with a high energy setting at a train of 30 s sonication with 30 s interval for 15 cycles using a Bioruptor Sonicator. After removing the insoluble debris, the cell lysate was pre-cleared with 40 μL of protein A-agarose beads (Millipore) for 2 h at 4°C with slow rotations. The immunoprecipitated DNA were enriched as described above in ChIP-seq.

The immunoprecipitated DNA was dissolved in 20 μL of 10 mM Tris-HCl pH 7.5 and then sonicated with a high energy setting at a train of 30 s sonication with 30 s interval for 5 cycles using a Bioruptor Sonicator. Vazyme DNA library preparation kit was used to construct a HiChIP library with modifications. After end repair and DNA adaptor ligation steps, 10 μL of washed streptavidin beads were resuspended with 100 μL of 2 × Biotin Binding buffer (10 mM Tris-HCl pH 7.5, 1 mM EDTA and 2 M NaCl). The streptavidin beads were added to the ligated DNA and rotated for 15 min at room temperature to enrich biotin-labeled DNA. The captured beads were washed twice with 1 × Tween Washing buffer (5 mM Tris-HCl pH 7.5, 0.5 mM EDTA, 1 M NaCl and 0.05% Tween 20). The washed beads were resuspended in 20 μL of H_2_O and amplified by PCR (95°C, 3 min; 98°C, 20 s, 60°C, 15 s, 72°C, 30 s for 13 cycles; and a final extension at 72°C, 5 min). After DNA purification, all HiChIP libraries were sequenced on an Illumina platform.

#### 4C

4C experiments were performed as previously described^16^ with slight modification. Briefly, ∼2 million cells were fixed with 2% formaldehyde and permeabilized with ice-cold permeabilization buffer (50 mM Tris–HCl pH 7.5, 150 mM NaCl, 5 mM EDTA, 0.5% NP-40, 1% Triton X-100, and 1 × protease inhibitors). The permeabilized cells were digested with Dpn II (NEB R0543S). The digested DNA fragments were ligated with T4 DNA ligase (NEB M0202S) and de-crosslinked. The ligated DNA was purified by phenol/chloroform extraction and ethanol precipitation, and sonicated to 200-600 bp fragments. Targeted fragments were linearly amplified using a 5’ biotin-labeled primer specific to the anchor region (Supplementary Table 2). The amplified ssDNA was enriched with Streptavidin (ThermoFisher 11206D), ligated with adapters (Adaptor-upper and Adaptor-lower, Supplementary Table 2), and amplified by PCR with anchor-specific P5-forward primers (Supplementary Table 2) and indexed P7-reverse primers. The amplified libraries were sequenced on an Illumina platform.

#### In situ Hi-C

*In situ* Hi-C experiments were performed as previously described^10^ with slight modification. Briefly, for each Hi-C experiment, 5 million cells were cross-linked with 1% formaldehyde at room temperature for 10 min, quenched with 0.125 M glycine, and lysed with 250 μL of ice-cold lysis buffer (10 mM Tris-HCl pH8.0, 10 mM NaCl, 0.2% NP-40) twice to obtain nuclei. The nuclei were then incubated with 50 μL 0.05% of SDS at 62°C for 10 min, quenched with Triton X-100, and digested with 100 units Mbo I (NEB R0147M) in a 250 μL volume overnight at 37°C. After heat inactivation of Mbo I at 62°C for 20 min, The Mbo I DNA ends were filled and labelled with biotin by adding 50 μL fill-in mix containing biotin-14-dATP (Thermo 19524016), dCTP, dTTP, dGTP, and DNA Polymerase I, Large (Klenow) Fragment (NEB M0210L) and incubation at 37°C for 1 h.

After proximity ligation at room temperature for 4 h using T4 DNA ligase (NEB M0202S), the samples were reversely cross-linked at 68°C overnight. DNA was precipitated with ethanol and sonicated for fragmentation. 300-500 bp DNA fragments were selected using AMPure XP beads (Beckman A63881). The streptavidin beads (Thermo 11206D) were then used to enrich the biotin-labelled DNA fragments. The on-bead biotin-DNA was washed stringently for a train of twice 2-min vortices at 55°C and used for library construction. NEB end-repair module (NEB E6050S) was used to blunt DNA ends and Klenow exo^-^ (NEB M0212S) was used to add an “A” at the 3’ ends. Illumina U-type adaptors were ligated to both DNA ends. The ligated adaptors were cleaved with the USER enzyme (NEB M5505S). After 10-12 cycles of PCR amplification with Illumina P5/P7 primers, the DNA products were purified with the AMPure XP beads (Beckman A63881) and paired-end sequenced on an Illumina NovaSeq platform.

#### Data analysis of ChIP-nexus

Raw reads were trimmed using Cutadapt (v2.10)^61^ to remove the first 10 bp barcode and adaptor sequences. All reads at least 20 bp in length after trimming were aligned to the human reference genome GRCh37/hg19 using Bowtie2 (v2.3.5.1)^53^. Reads from repeated samples were merged, sorted, and indexed using Samtools (v1.12)^56^. Narrow peaks were called using MACS2 (v2.2.7.1)^57^ with a q-value threshold of 0.001. Bedtools (v2.30.0)^55^ was used to select overlapping peaks and the R package Vennerable (v3.1.0.9000) was used to generate Venn diagrams. The genomecov function of Bedtools was used to calculate the coverage of the mapped reads. The summits of read coverage in peaks around the forward or reverse CTCF motifs were collected to generate violin plots and boxplots with ggplot2.

Read counts were normalized to reads per kilobase per million mapped (RPKM) using the bamCoverage module of Deeptools (v3.5.1)^41^ with a bin size of 20 bp, and converted to bedGraph format for visualization in the UCSC genome browser. Heatmaps were generated using the plotHeatmap module of Deeptools. Scatter plots were generated by a Python script with peak read counts calculated by Bedtools^55^. The coverage of the 5’ ends of the positive or negative strand was calculated using the genomecov module of Bedtools. Footprint density profiles of CTCF, ZNF143, and RAD21 for the forward and reverse CBS elements were generated using ggplot2.

#### DNA motif analysis

The MEME suite (v4.12.0)^62^ was used for DNA motif analysis. For the SBS element, ZNF143 ChIP-nexus narrow peaks not overlapped with CTCF peaks were used for motif searching. The 1 kb upstream regions of peaks were used to establish a background model using fasta-get-markov. Two thousand random peaks were used for motif finding with the following parameters: -revcomp -w 20 -mod zoops -env 0.0001. The top motif was used as the core motif. The 20 bp-regions flanking the core motif were used for motif finding again with a threshold of 0.001. Motifs were combined together to form different types of motifs using the SpaMo module of MEME. The MEME FIMO module was used to find the binding positions and orientations of ZNF143 binding sites. All motif types were scanned sequentially in all peaks. Once a motif was found, the peak was excluded from subsequent scans.

### Quantification and statistical analysis

#### Data analysis of ChIP-seq

Reads from ChIP-seq were mapped to the human reference genome GRCh37/hg19 using Bowtie2. Peaks were called using MACS2^57^ with default parameters. All mapped SAM files were sorted and indexed using Samtools^56^ and normalized to RPKM using the bamCoverage module of Deeptools. The bedGraph file generated from Deeptools was uploaded to the USCS genome browser for visualization.

#### Data analysis of RNA-seq

RNA-seq raw FASTQ files were aligned to the GRCh37/hg19 reference genome using STAR (v2.7.3a)^63^ with default parameters. The BAM files were analyzed using Cufflinks (v2.2.1)^58^ to calculate expression levels of transcripts in fragments per kilobase of exon per million fragments mapped (FPKM). The raw counts were used to identify differential expression genes using DESeq2^60^ with parameters of abs(log2FoldChange) > 1 and pvalue < 0.05. The volcano plot that displays differential expressed genes was generated by ggplot2. BAM files were converted to the bedGraph format using Deeptools for visualization in the UCSC genome browser.

#### Data analysis of 4C

4C FASTQ raw reads were aligned to GRCh37/hg19 using Bowtie2 with default parameters. Reads per million (RPM) interaction values were calculated using the r3Cseq program (v1.38.0)^59^. The generated bedGraph files were used for visualization in the UCSC genome browser. The interaction files from r3Cseq were used to calculate the interaction differences.

#### Data analysis of Hi-C

Raw reads were mapped to GRCh37/hg19 to generate contact maps using HiC-Pro (v3.0.0)^54^. The produced allValidPairs file was transformed to a hic format file using the hicpro2juicebox script from HiC-Pro for further analyses with the Juicer tools^64^. The hic file was converted to the cool format using hic2cool (v0.8.3). The Hi-C contact matrix was balanced using the Knight-Ruiz (KR) method and normalized to a depth of 100 million contacts.

Loops with 1D genomic distance of more than 4 kb were detected using the detect function of Chromosight (v1.6.3)^68^ with the parameters --min-dist 40,000 -p 1e-5. Aggregate peak analysis (APA) for loops was performed using cooltools(v0.5.2)^66^ and coolpup.py (v1.0.0)^67^.

TAD domains were called using the HMM (hidden Markov models) method as previously described^70^. Briefly, the sparse matrix was transformed into a dense matrix, which was used to calculate the directionality index (DI) score with a bin size of 10 kb and a window size of 2 Mb. It is assumed that each bin has a hidden state which marks the upstream boundary of TADs, the downstream boundary of TADs, or not a boundary. The DI score was used to predict the hidden states of all bins using hidden Markov models. Consecutive bins with the same state were merged into regions. We filtered out regions composed of less than 3 bins or with a median posterior bin probability lower than 0.99. If the upstream and downstream regions of a filtered region had the same bin state, then the filtered region transforms all bins to that state. Otherwise, all bins were assumed to be non-boundary ones. Finally, TADs are defined as the area both downstream to an upstream boundary region and upstream to a downstream boundary region. Aggregate domain analysis (ADA) for TADs was calculated using a custom Python script.

The DI score was converted to bedGraph format for visualization in the UCSC genome browser. The aggregated DI values around TAD boundaries were calculated using a custom Python script.

Hi-C *cis-*Eigenvector 1 values and Pearson’s correlation matrix were computed using the eigenvector and Pearson’s function of Juicer, respectively, at 100-kb resolution. The *cis-*Eigenvector 1 values were converted to bedGraph format for visualization in the UCSC genome browser. The density scatterplot and Pearson’s R between different samples were calculated using a custom Python script. Virtual-4C profiles were generated using Fan-C^65^ at 10-kb resolution.

Insulation scores were called as previously described^71^. Briefly, the sparse matrix was transformed into a Dekker format dense matrix. For all bins at 10 kb resolution, we calculated the mean interaction strength between the upstream 50 bins and downstream 50 bins as the insulation score. The aggregated insulation scores around TAD boundaries were calculated using a custom Python script.

#### Data analysis of HiChIP

HiChIP raw data were aligned to GRCh37/hg19 to generate contact maps using the HiC-Pro pipeline. The aligned reads from HiC-Pro and the peaks from ChIP-nexus data were used to call loops with the hichipper software (v0.7.7)^42^. Loops with less than 20 kb in length or with fewer than 2 paired-end tags were filtered out.

Loops with both anchors overlapped with ZNF143 peaks but not CTCF peaks were defined as the SBS loops. Loops with both anchors overlapped with CTCF peaks but not ZNF143 peaks were defined as the CBS loops. Loops with one anchor overlapped with CTCF peak and the other anchor overlapped with ZNF143 peaks were defined as the SBS-CBS loops.

Loops with both anchors overlapped with promoters were defined as the promoter-promoter (P-P) loops. Loops with both anchors overlapped with enhancers were defined as the enhancer-enhancer (E-E) loops. Loops with one anchor overlapped with a promoter and the other anchor overlapped with an enhancer were defined as the promoter-enhancer (P-E) loops.

The loop data were converted into the interacting format for visualization as arcs in the UCSC Genome Browser or the WashU Epigenome Browser. To validate the SBS-SBS, SBS-CBS, and CBS-CBS loops generated by HiChIP in Hi-C data, the HiChIP loop anchors were refined to the closest Mbo I restriction fragments and further extended 12 fragments on both sides. The counts of Hi-C valid pairs generated by HiC-Pro in each loop were normalized by the total valid-pair counts and then multiplying by 100 million. The *P*-value was calculated using a paired Student’s *t*-test.

#### Statistical analysis

ChIP-nexus, HiChIP, Hi-C, RNA-seq, ChIP-seq, and 4C experiments were performed with at least two biological replicates. All statistical tests were calculated using R and python scripts. Data were expressed as mean ± standard error of the mean (SEM) or confidence interval (CI). Statistical significance values were calculated using an unpaired Student’s *t*-test. *P* ≤ 0.05 was indicated as ‘*’. *P* ≤ 0.01 or 0.001 was indicated as ‘**’ or ‘***’, respectively.

## Data and code availability

λ High-throughput sequencing data and processed files have been deposited in the Gene Expression Omnibus (GEO) database under accession number GSE236637.
λ This paper does not report original code.
λ Any additional information required to reanalyze the data reported in this paper is available from the lead contact upon request.

## Supplemental information

Supplementary Figures

Supplementary Tables

**Table S1.** SBS loops identified by ZNF143 HiChIP, Related to Figure 4

**Table S2.** Oligonucleotides used in this study, Related to Figures 1, 4-7

## Key Resources Table

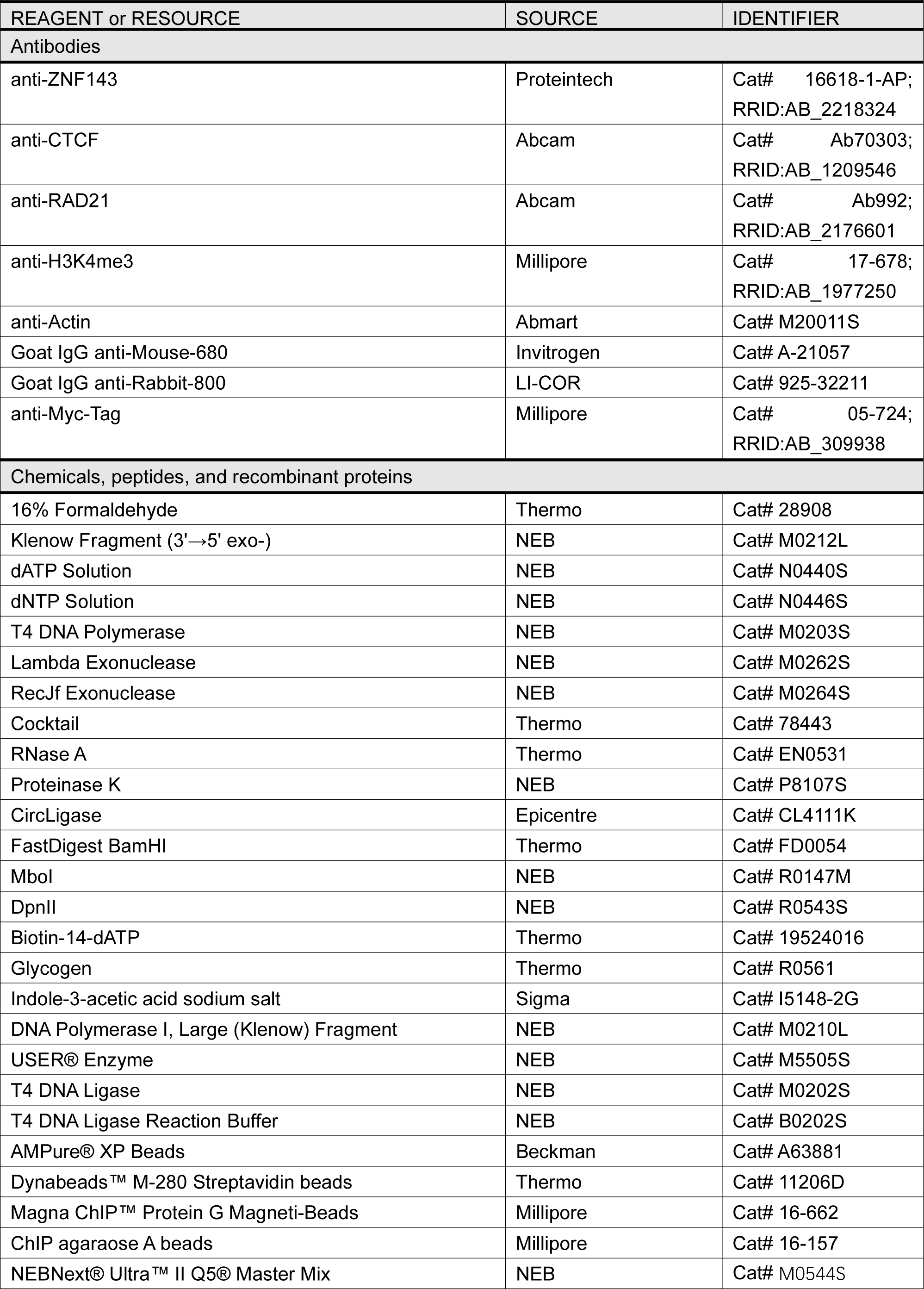
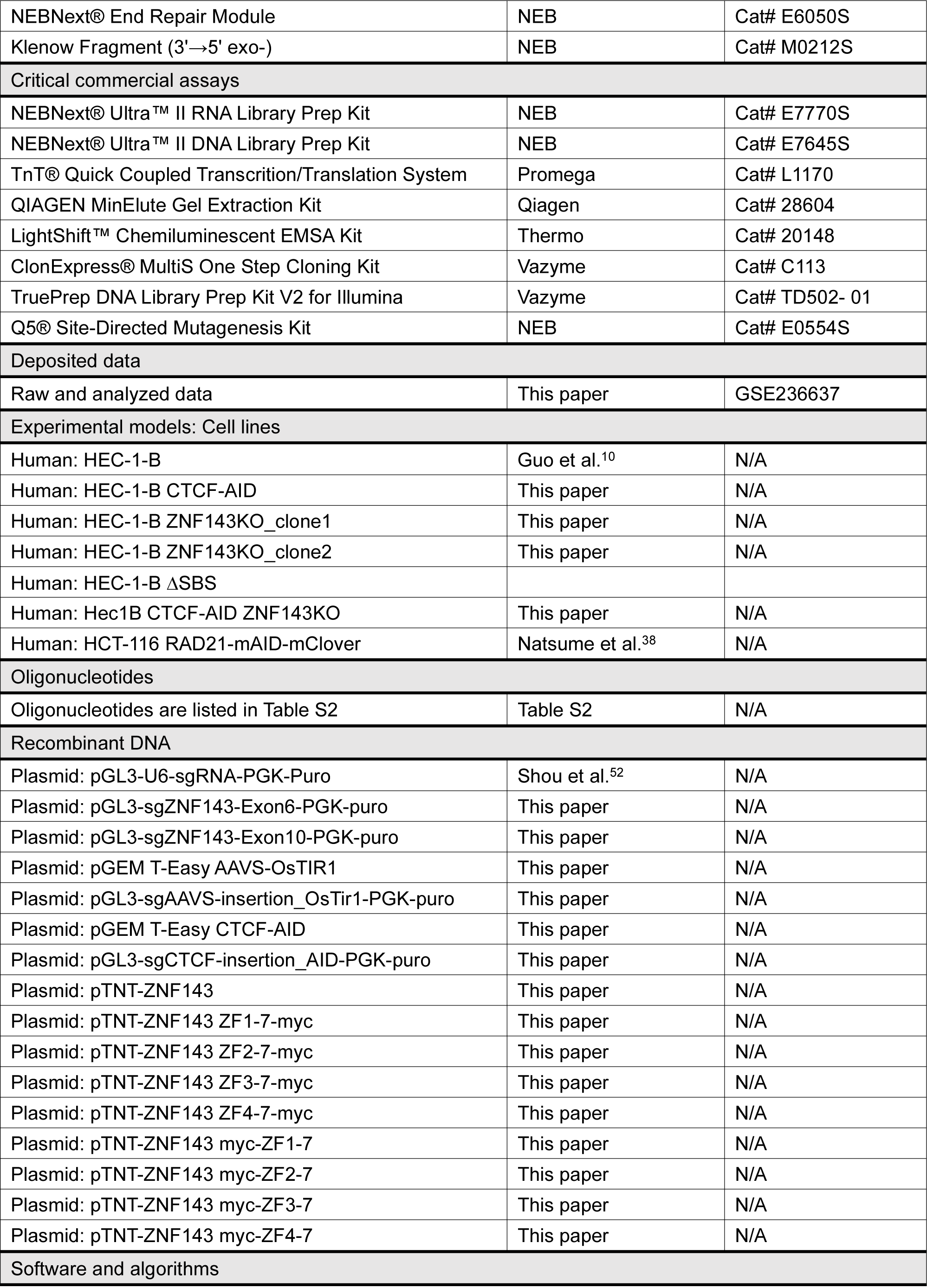
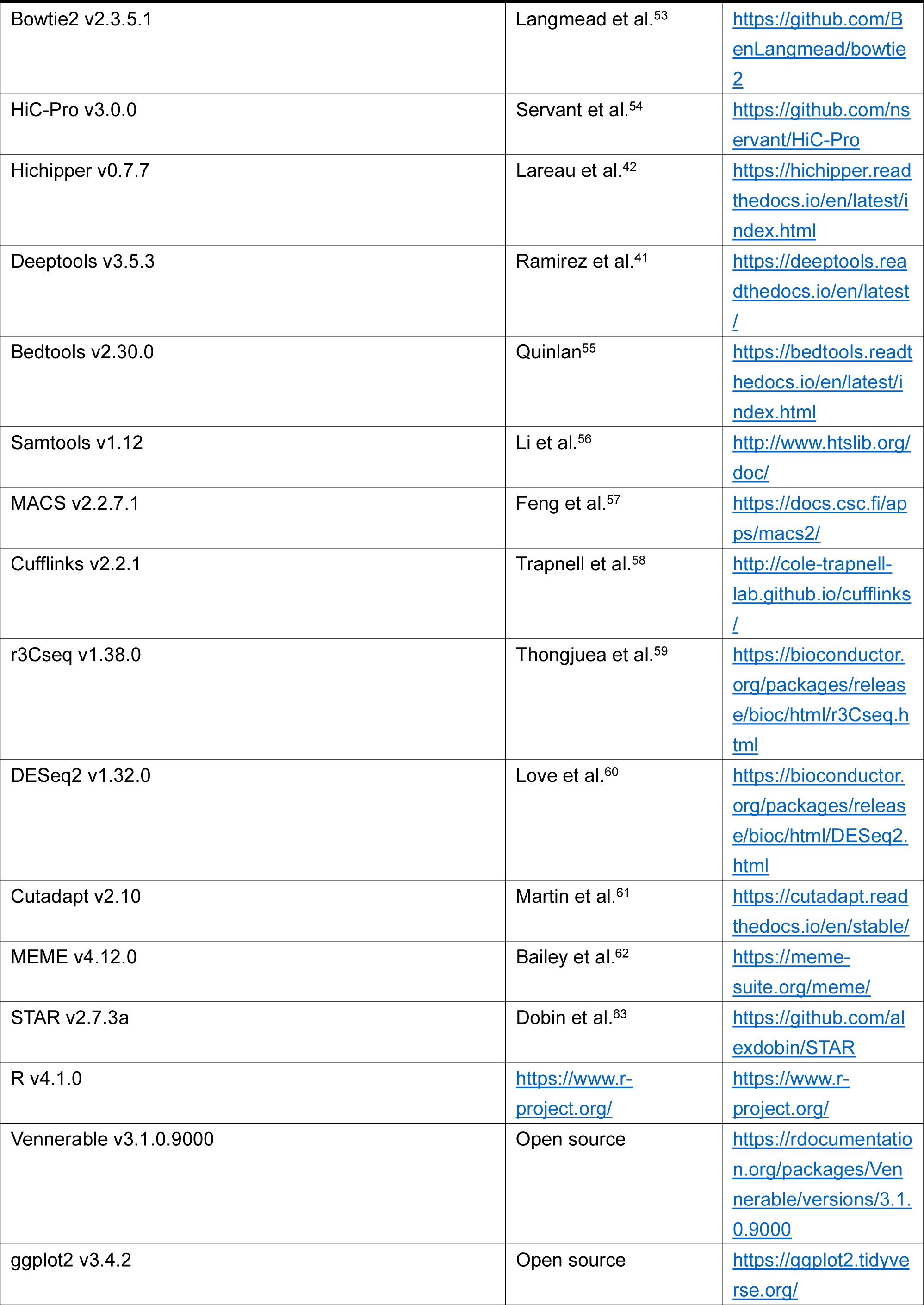
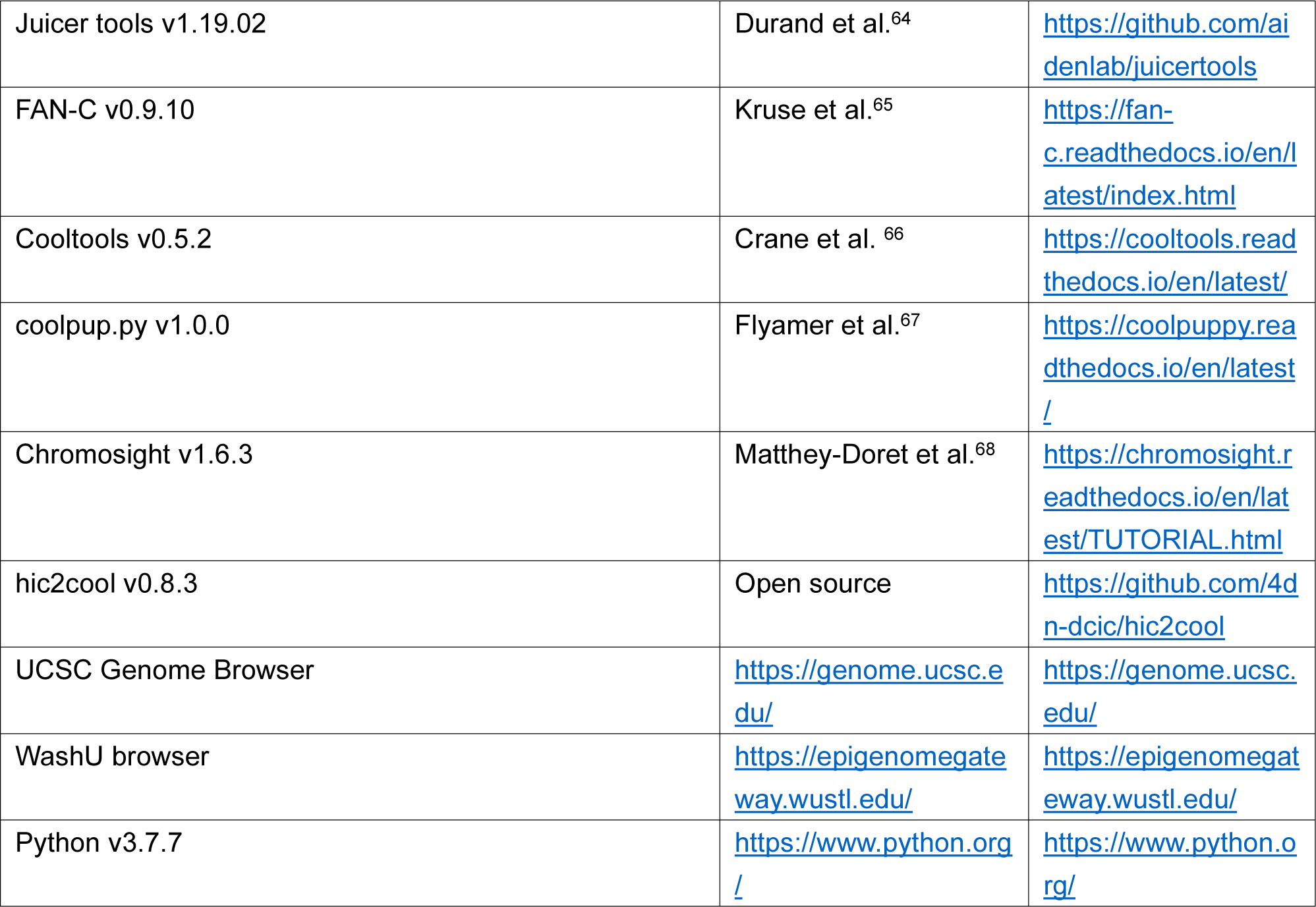

---

**Figure S1.**
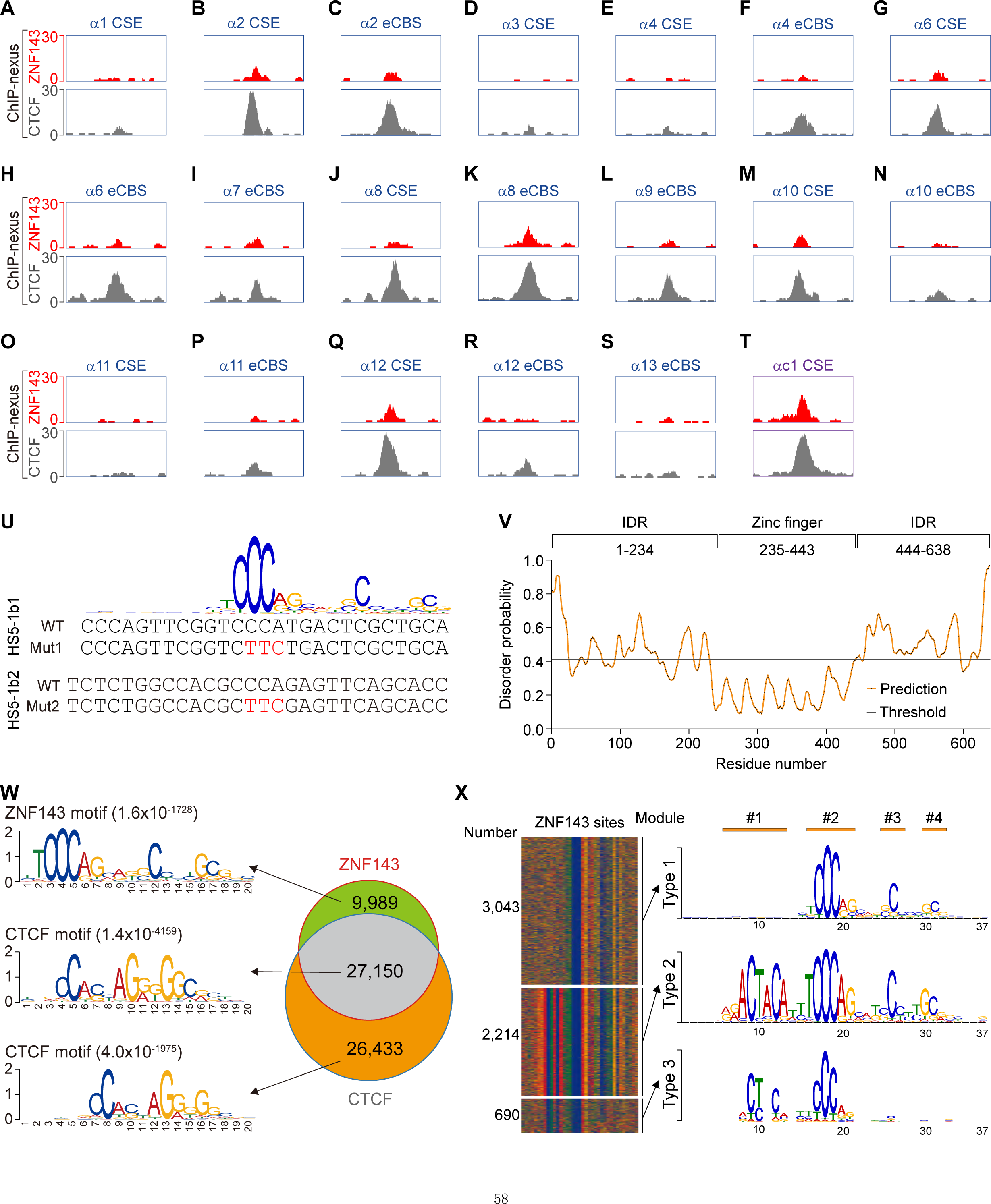
Directional ZNF143 recognition of SBS elements, Related to Figure 1. (**A**-**T**) ZNF143 and CTCF ChIP-nexus profiles in HEC-1-B cells showing the colocalization of ZNF143 and CTCF at the *CSE* and *eCBS* elements of human *PCDHα* variable promoters. CSE, conserved sequence element; eCBS, exonic CBS. (U) Sequences of the WT and mutant (Mut1 and Mut2) probes of *HS5-1b1* and *HS5-1b2* SBS elements. (V) Probability of the disordered amino acid sequences of ZNF143. IDR, intrinsically disordered region. (W) Venn diagram of ZNF143 and CTCF sites as well as their motifs. (X) ChIP-nexus heatmap and motifs of three types of ZNF143 sites. Motifs were produced by the MEME software.

**Figure S2.**
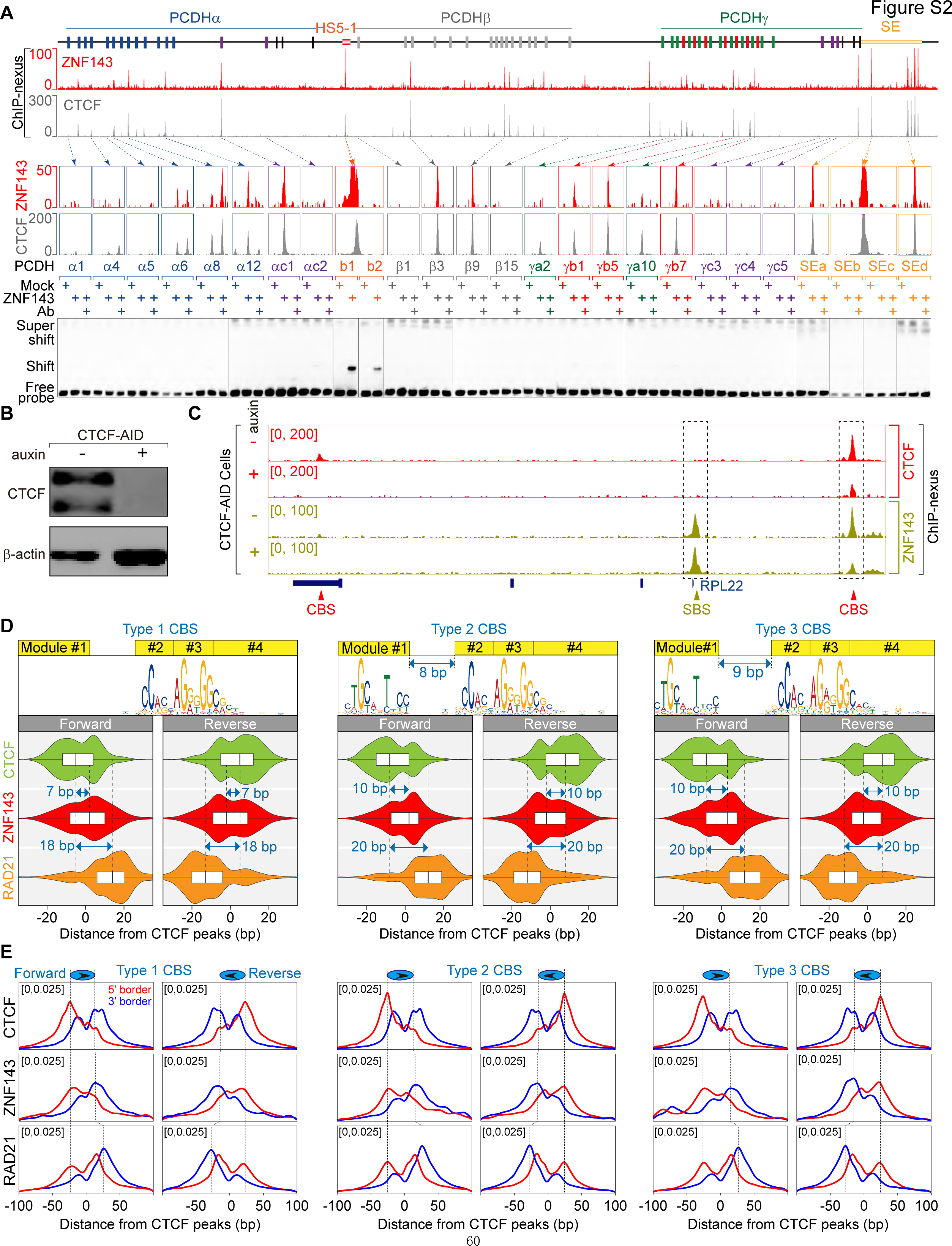
CTCF recruits ZNF143 to the CBS elements in vivo, Related to Figure 2. (A) ZNF143 and CTCF ChIP-nexus profiles of the *PCDH* clusters with corresponding EMSA below, suggesting indirect binding of ZNF143 to CBS elements and direct binding to the two SBS elements (*HS5-1b1* and *HS5-1b2*). (B) Western blot of CTCF in CTCF-AID cells treated with or without auxin, confirming CTCF depletion after auxin induction. (C) ZNF143 and CTCF ChIP-nexus profiles of the *RPL22* locus in CTCF-AID cells, showing decreased enrichment of ZNF143 at CBS, but not SBS, elements upon CTCF degradation. (D) Violin plots of the distributions of ZNF143, CTCF, and RAD21 ChIP-nexus peak summits of three types of forward or reverse CBS elements, showing the localization of ZNF143 between CTCF and RAD21. Type 1 CBS motif lacks Module1. Type 2 or 3 CBS motif contains Module1 at 8 or 9 bp upstream of Module2, respectively. (E) ChIP-nexus footprint profiles of ZNF143, CTCF, and RAD21 of three types of forward or reverse CBS elements. Arrows indicate CBS orientations. Red or blue lines show 5’ or 3’ borders of footprints.

**Figure S3.**
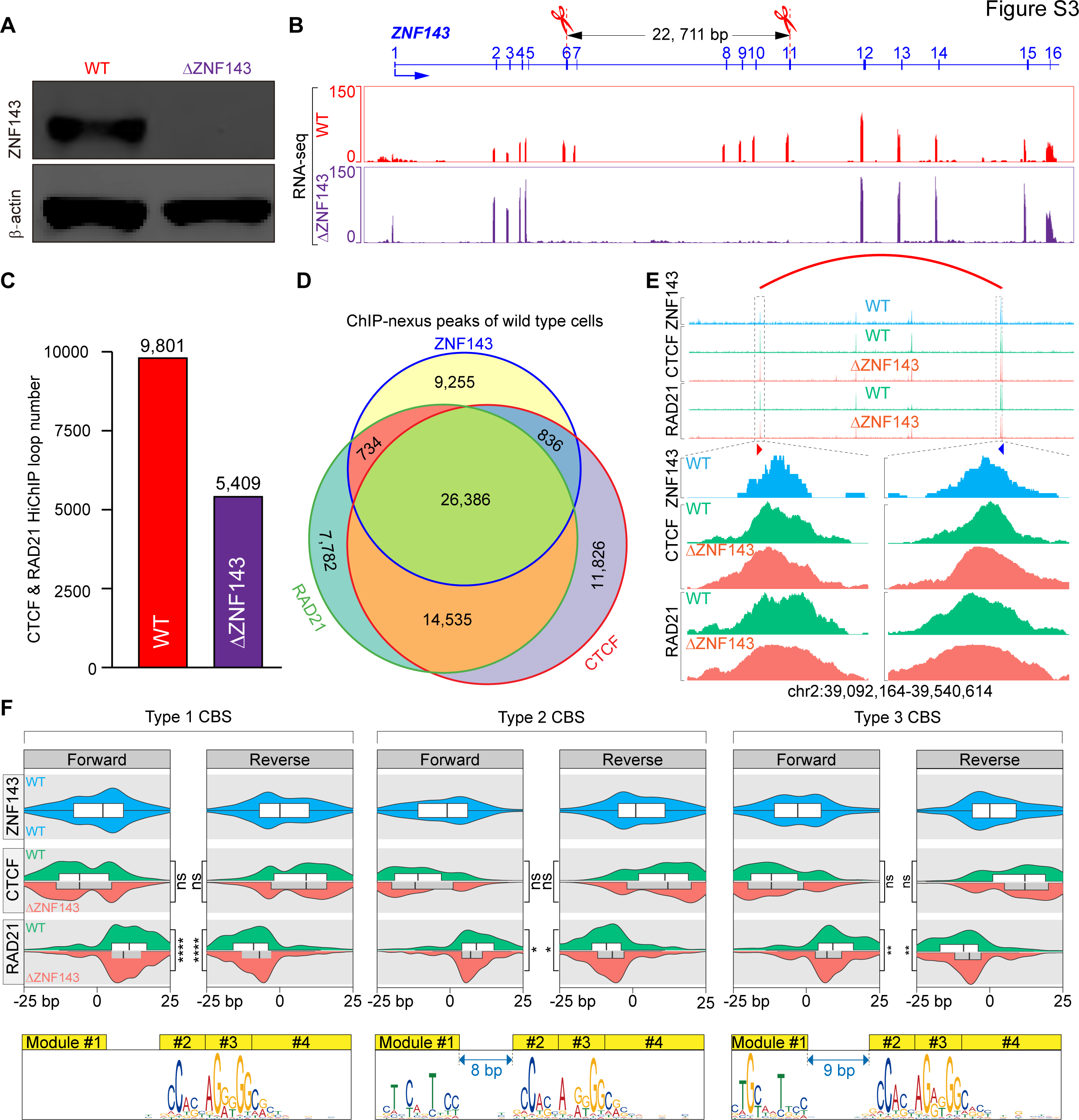
ZNF143 deletion compromises RAD21 but not CTCF enrichments, Related to Figure 3. (A) Western blot confirming ZNF143 deletion. (B) RNA-seq profiles of the *ZNF143* in WT and DZNF143 cells. (C) Number of CTCF and RAD21 HiChIP loops in WT and DZNF143 cells. (D) Venn diagram of ZNF143, CTCF, and RAD21 ChIP-nexus peaks. (E) A shift of the RAD21 ChIP-nexus peaks toward CTCF at loop anchors upon ZNF143 deletion. Red or blue arrow indicates the forward or reverse CBS elements. (F) Violin plots of ZNF143, CTCF, and RAD21 ChIP-nexus peak summits of three types of forward or reverse CBS elements in WT and DZNF143 cells, showing a closer proximity between RAD21 and CTCF upon ZNF143 deletion.

**Figure S4.**
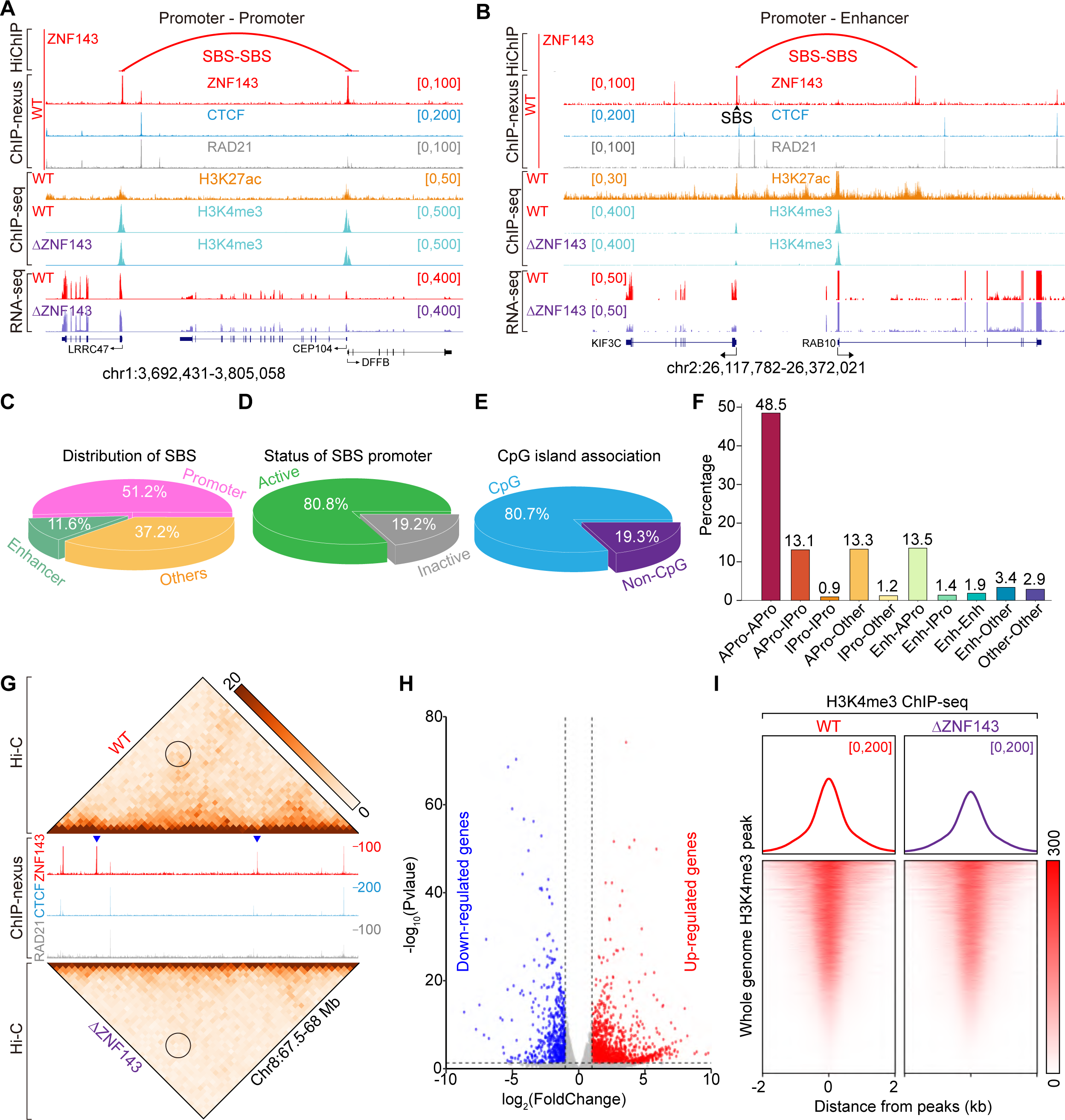
ZNF143 is required for SBS-SBS loop formation, Related to Figure 4. (**A and B**) SBS loops between promoters (**A**) or between a promoter and enhancer (**B**). (C) Localizations of SBS elements at the promoter, enhancer, and other regions. (D) Transcriptional status of SBS promoters. (E) CpG island association of SBS promoters. (F) Distribution of SBS loops with different combinations of *cis*-regulatory elements at both anchors, showing that most SBS loops are anchored at active promoters at least on one side. APro: active promoter, IPro: inactive promoter, Enh: enhancer. (G) Hi-C contact maps showing decreased interactions between two SBS loop anchors upon ZNF143 deletion. Bin size, 10 kb. (H) Volcano plots showing differentially expressed genes in DZNF143 and WT cells. (I) Decreased enrichments of H3K4me3 upon ZNF143 deletion.

**Figure S5.**
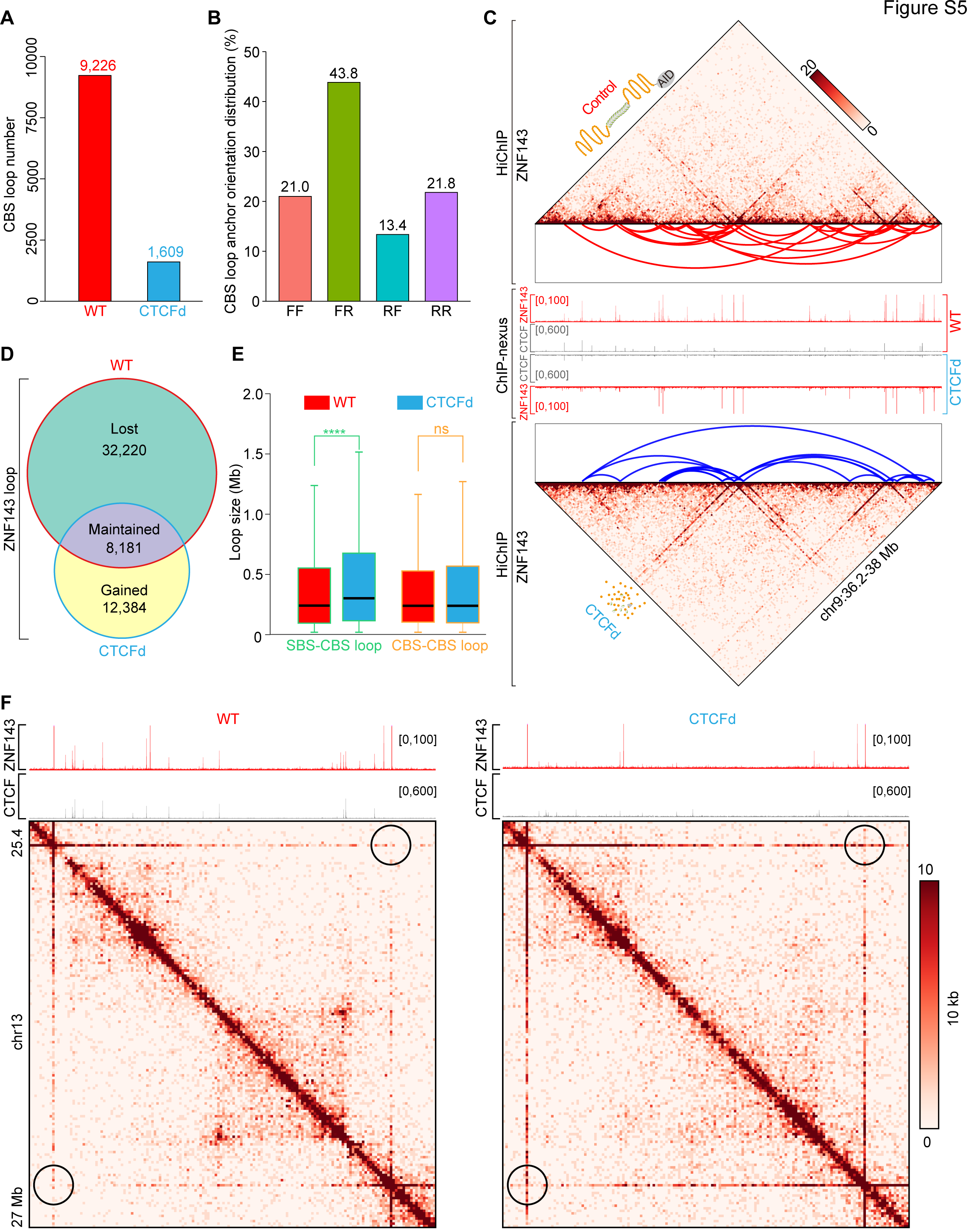
Aberrant formation of large-sized SBS loops upon CTCF degradation, Related to Figure 5. (A) CBS loop number in auxin-treated (CTCFd) or untreated (WT) CTCF-AID cells. (B) Distribution of CBS loop orientation revealed by ZNF143 HiChIP. (C) ZNF143 HiChIP heatmap revealing the formation of large chromatin loops upon CTCF degradation. Bin size, 10 kb. (D) Number of ZNF143 HiChIP CBS loops that are lost, maintained, or gained after CTCF degradation. (E) Distributions of SBS-CBS and CBS-CBS loops showing an increase of the average size of SBS-CBS, but not CBS-CBS loops, upon CTCF depletion. (F) ZNF143 HiChIP heatmap showing that a large SBS loop is strengthened upon acute CTCF degradation.

**Figure S6.**
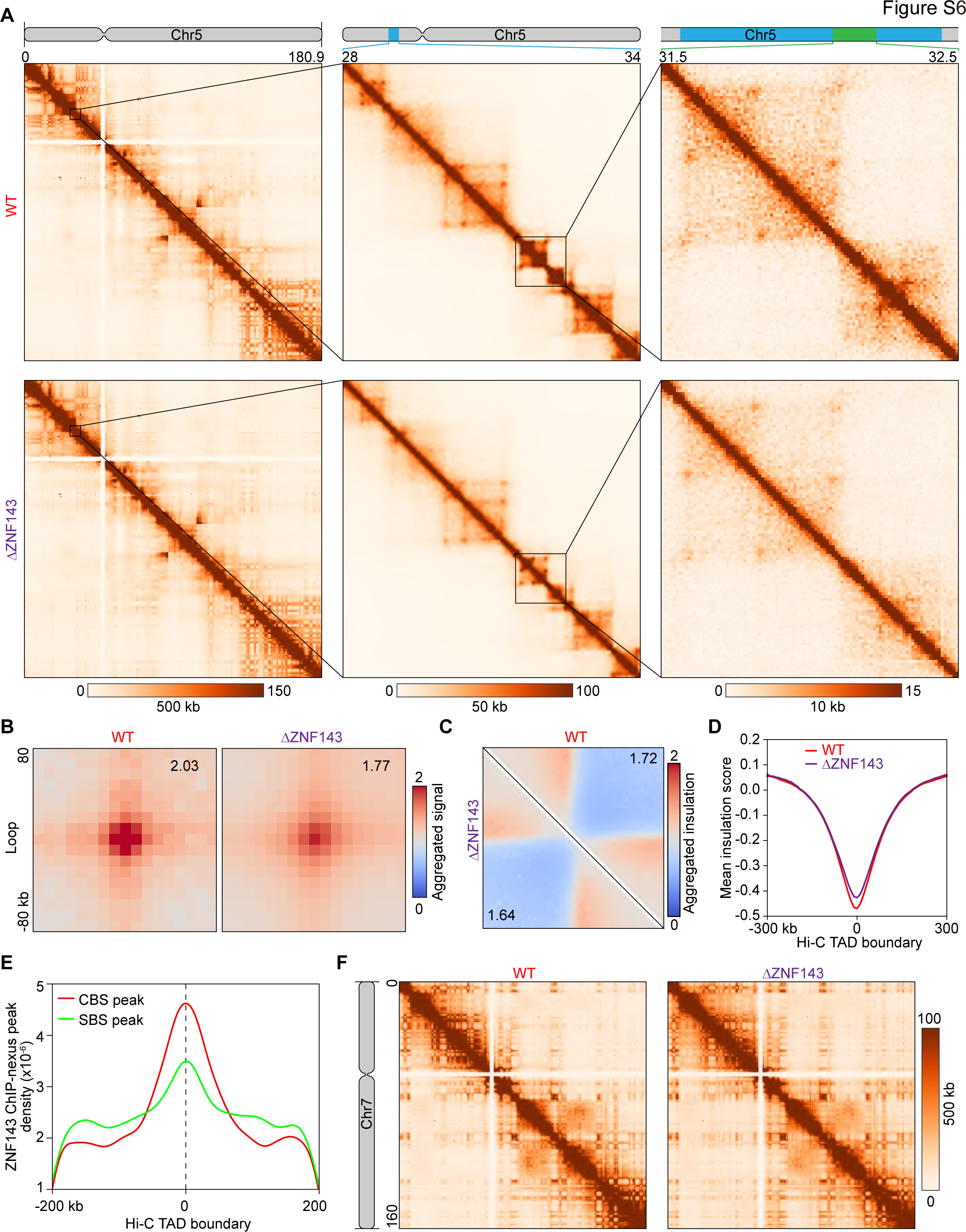
ZNF143 deletion weakens TAD boundaries, Related to Figure 6. (**A**) Hi-C contact matrices of chromosome 5: the whole chromosome, at 500 kb resolution (left); 28-34 Mb/50 kb resolution (middle); 31.5-32.5 Mb/5 kb resolution (right), showing decreased chromatin contacts upon ZNF143 deletion. The 1D region corresponding to a contact matrix is indicated in the diagrams above. (B) APA plot for Hi-C loops called from WT and DZNF143 cells, showing global weakening of chromatin loops upon ZNF143 deletion. The APA score is indicated in the upper-right corner of each panel. (C) Heatmap of insulation scores of TAD boundaries in WT and DZNF143 cells. (D) Averaged insulation scores in WT and DZNF143 cells. (E) Distribution of SBS or CBS peaks in 200-kb regions flanking Hi-C TAD boundaries. (F) Hi-C contact maps at 500 kb resolution across the entire chromosome 7.

**Figure S7.**
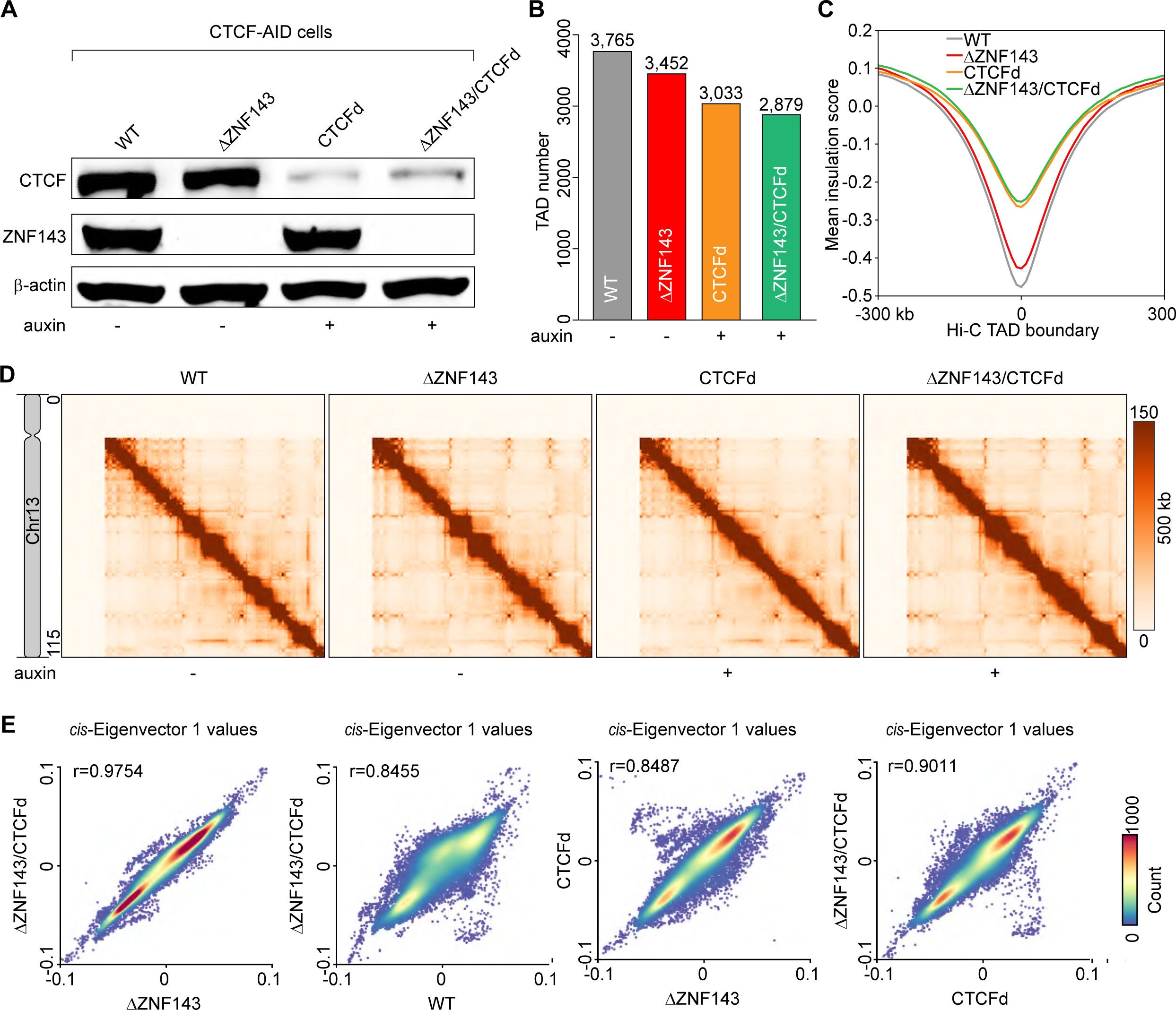
ZNF143 and CTCF collaborate to organize 3D genome, Related to Figure 7. (A) Western blot of WT control, ZNF143 deletion (DZNF143), acute CTCF degradation (CTCFd), and ZNF143 deletion combined with CTCF degradation (DZNF143/CTCFd). (B) Bar graph showing the further decrease of TAD number upon ZNF143 deletion combined with CTCF degradation. (C) Averaged insulation scores of all Hi-C TAD boundaries showing a global decrease in TAD boundary insulation upon ZNF143 deletion and CTCF degradation. (D) Hi-C contact maps at 500 kb resolution across the entire chromosome 13. (E) Scatter plot of the Pearson’s correlation of *cis-*Eigenvector 1 values showing distinction between ZNF143 deletion and CTCF degradation. This suggests that ZNF143 deletion, but not CTCF degradation, alters genome compartmentalization.

